# Spatial Transcriptomics Using Multiplexed Deterministic Barcoding in Tissue

**DOI:** 10.1101/2022.08.30.505834

**Authors:** Johannes Wirth, Nina Compera, Kelvin Yin, Sophie Brood, Simon Chang, Celia P. Martinez-Jimenez, Matthias Meier

## Abstract

In this study, we present a multiplexed version of deterministic barcoding in tissue (xDbit) to acquire spatially resolved transcriptomes of nine tissue sections in parallel. New microfluidic chips were developed to spatially encode mRNAs over a total tissue area of 1.17 cm^2^ with spots of 50 μm×50 μm. Optimization of the biochemical protocol increased read and gene counts per spot by one order of magnitude compared with previous reports. Furthermore, the introduction of alignment markers allows seamless registration of images and spatial transcriptomic spot coordinates. Together with technological advances, we provide an open-source computational pipeline to transform raw sequencing data from xDbit experiments into the AnnData format. The functionality of xDbit was demonstrated by the acquisition of 18 spatially resolved transcriptomic datasets from five different murine organs, including cerebellum, liver, kidney, spleen, and heart. Factor analysis and deconvolution of xDbit spatial transcriptomes allowed for in-depth characterization of the murine kidney.

## Introduction

Single-cell transcriptomics (scT) has revolutionized the concept of cellular heterogeneity and led to the development of comprehensive reference maps of cells typically isolated from biopsies, tissues, and whole organisms (Macosko et al., 2015; Picelli et al., 2013; Rosenberg et al., 2018; Vitak et al., 2017). These methods elucidate cell-to-cell communication, and tissue architecture which play key roles in tissue homeostasis, tissue repair, and disease progression. However, tissue dissociation protocols cause loss of spatial information and alteration of cell type proportions, removing critical information to understand cellular crosstalk and the microenvironment. To overcome this limitation, spatial transcriptomics (ST) has been developed based on imaging, sequencing, or both methodologies (Moses and Pachter, 2022). Imaging-based methods exploit *in situ* hybridization probes to detect single transcripts with high spatial resolution down to the subcellular scale; however, the need for targeted probes limits the study to a predetermined set of genes (Chen et al., 2015; Eng et al., 2019). Instead, for sequencing-based techniques, RNAs are barcoded with DNA molecules to encode the spatial position. Enabling, untargeted detection of mRNAs of the whole transcriptome. While commercially available technology resolves tissue spots with diameters of the order of tens of microns, recent technical improvements made in high-definition spatial transcriptomics (HDST) (Vickovic et al., 2019) and STEREO-seq (Wu et al., 2021a) resolve transcripts down to spot sizes of sub-microns. Alternative methods, including sci-SPACE (Srivatsan et al., 2021) and XYZeq (Lee et al., 2021) barcode biomolecules within tissue spots before retrieval. In the proceeding step, nuclei or whole cells are isolated from the tissue, and their transcriptomes are sequenced next to positional barcodes. To select the most suitable ST method for an experiment, several parameters, namely spatial resolution, detection limit, screening area, accessibility, compatibility with existing workflows, and costs, are weighed against each other depending on the research question. For instance, high-resolution methods either require specialized equipment to manufacture the components and establish the analysis in the lab, or are proprietary, which leads to higher costs. Lower resolutions are, in turn, associated with the loss of single-cell resolution because of the larger resolved spot sizes. Larger spot sizes can be compensated for by integrating scT with ST and inferring the cell type composition of each spot (Andersson et al., 2020; Kleshchevnikov et al., 2022; Lopez et al., 2022; Ma and Zhou, 2022).

Deterministic barcoding in tissue (DBiT-seq) is a cost-effective and openly accessible platform to scale ST (Liu et al., 2020). DBiT-seq uses microfluidic channels to barcode tissue sections using DNA oligonucleotides and allows the integration of multi-omics information, including antibodies (Liu et al., 2022), epigenomics (Deng et al., 2022a), and chromatin accessibility readouts (Deng et al., 2022b).

In this study, we present multiple<u>x</u>ed <u>d</u>eterministic <u>b</u>arcoding in tissue (xDbit), a method for acquiring spatially resolved transcriptomes from nine fixed tissue sections in parallel. Optimization of the chemical protocol and workflow of the DBiT-seq method led to an increase in transcript reads and gene counts per 50 × 50 μm spot. The introduction of alignment marks onto the tissue sections enabled the seamless acquisition of transcriptomic reads and spatial registration with high-resolution images. Together with technological advances, we provide an open-source computational pipeline to transform the raw sequencing data from an xDbit experiment into Scanpy-compatible data file formats (Virshup et al., 2021; Wolf et al., 2018).

To demonstrate the functionality of xDbit, we acquired spatially resolved transcriptomic datasets of 18 tissue sections from five different murine organs, including the cerebellum, liver, kidney, spleen, and heart. Using the kidney as model tissue, we show that xDbit can be used in conjunction with factor analysis to perform an in-depth characterization of organs and identify spatially patterned genes. Finally, we demonstrated that xDbit can resolve rare cell types upon cell-type deconvolution using scT data, allowing cost-efficient research projects on spatiotemporal expression dynamics in longitudinal studies and multi-organ comparisons.

## Results

### Multiplexed Deterministic Barcoding in Tissue (xDbit)

To enable multiplexing, increase sequencing depth, and improve the imaging quality of the Dbit-seq methodology, we developed a multiplexed deterministic barcoding in tissue (xDbit) workflow (**Fig. 1A**). For an xDbit experiment, nine fresh frozen tissue sections with a maximal area of 0.4 × 0.4 mm were positioned in a 3 × 3 grid layout on a glass substrate (**Figure S1H**). Tissue sections were fixed with PFA and the nuclei, cytoskeleton, and selected proteins were stained using a standard immunofluorescence protocol (Methods). High-resolution images were acquired before the xDbit run to obtain high-quality images without the introduction of artifacts from the downstream ST processing steps. Subsequently, mRNAs within tissue sections were reverse transcribed using a 3D printed 9-well chip (**Figure S1F**), which separated each section and reduced the reaction volume to 80 μL per sample. The reverse transcription (RevT) primer carried a hybridization site to ligate the spatial barcodes in the following working steps, and a poly(T) 3’-end to bind to and reverse transcribe all polyadenylated mRNAs (**Figure S2**). Analogous to DBiT-seq, spatial barcoding of the resulting cDNA was performed using two sequentially aligned polydimethylsiloxane (PDMS) chips. The first PDMS chip was clamped onto the tissue, creating 38 parallel and horizontally aligned microchannels (50 μm × 50 μm) on top of each tissue section, and allowing DNA barcodes to be flushed over the tissue. The DNA barcodes were ligated to the cDNA within the underlying tissue, thereby encoding the positions of the horizontally directed channels.

**Figure 1.**
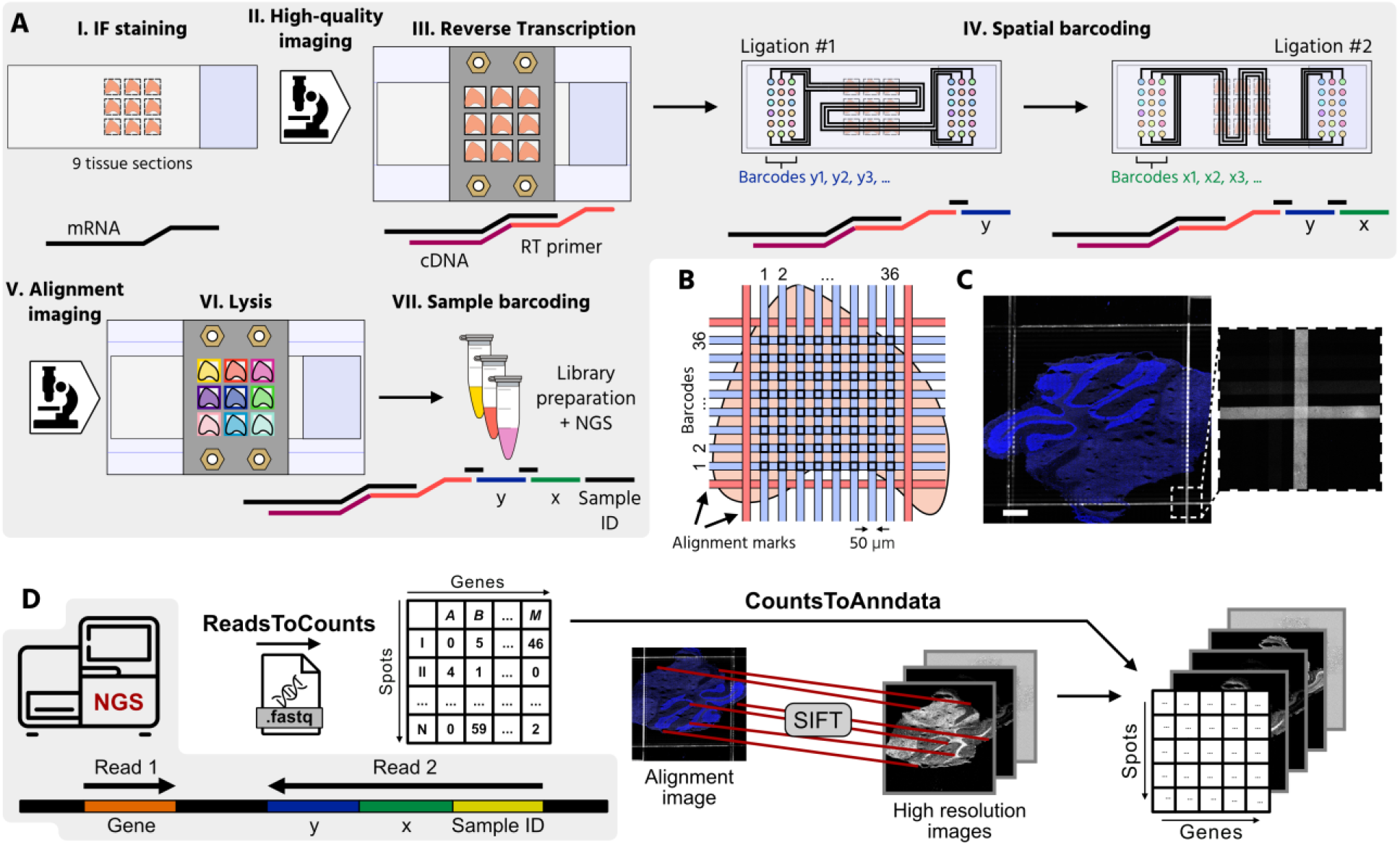
Multiplexed deterministic barcoding in tissue (xDbit). (**A**) Workflow of xDbit: (I) Deposition of PFA-fixed cryosections with a thickness of 10 μm onto a glass substrate in a 3 × 3 matrix layout. (II) Acquisition of high-quality immunofluorescence images after standard antibody staining. (III) Reverse transcription of mRNA into cDNA in all tissue sections with the help of a 3D-printed 9-well chip. (IV) Spatial barcoding of the cDNA in two sequential ligation steps. Each ligation step is performed with an individual PDMS chip. The two chips were designed with an orthogonal serpentine channel layout to generate a 38 × 38 spot array on the tissue sections. The channel intersections generate a spatially encoded spots with an area of 50 μm × 50 μm and a center-to-center spot distance of 100 μm. Alignment marks to register IF images and spatial transcriptomic coordinates are generated by filling the outermost channels with a fluorescently labelled anti-BSA antibody. The antibody targets the blocking reagent BSA, which was deposited on the tissue surface during the IF staining. (V) Re-imaging of the tissue section to obtain an alignment image. (VI) Lysis of the 9 individual tissues is achieved using the 3D printed 9-well chip. (VII) Each sample is individually indexed before sequencing. (**B**) Channel layout to generate alignment markers for the registration of high-resolution images and spatial transcriptomic spot coordinates. (**C**) Representative fluorescence images of the alignment marks with zoom-in image. Blue and white denote DAPI counterstain and anti-BSA staining, respectively. Scale bar: 500 μm. (**D**) Schematic of the xDbit computational pipeline. NGS reads are transformed into a spot-gene count matrix using the *ReadsToCounts* script. Alignment-and high-quality images are registered via their DAPI signal using the SIFT algorithm (Lowe, 2004) and then aligned to the count matrix with the *CountsToAnndata* script. The resulting integrated datasets are compatible with *Scanpy* and *Squidpy* analysis pipelines (Palla et al., 2022; Virshup et al., 2021; Wolf et al., 2018).

The second PDMS chip resembles the first chip, with the difference that the 38 microchannels run vertically over the tissue section to barcode the cDNA in the tissue via ligation with an identifier for the vertically directed channel. The spatial barcoding resulted in a grid of 1444 uniquely barcoded spots, each with a width of 50 μm. In contrast to the original DBiT-seq approach, 38 microchannels were guided in serpentines over the glass substrate, which allowed us to address nine tissue sections in parallel and increased the scanning area from 25 to 116.64 mm^2^ (4.66-fold increase). Importantly, we found that dehydration of the tissue sections with ethanol was essential to ensure the optimal attachment of the microfluidic chips. To enable registration of the spatial transcriptomic spots to the image data, the two outermost channels were filled with an alignment marker solution (**Fig. 1B**) consisting of an anti-BSA antibody that binds to the BSA-blocked surface. After the second round of ligation, the tissue sections were imaged again to record alignment marks and stained nuclei (**Fig. 1C**). Finally, a 9-well 3D-printed chip was attached to the slide to lyse the tissue sections individually. Within xDbit, tissue multiplexing can be achieved after either reverse transcription with barcoded primers or sample retrieval by indexed library preparation.

For the analysis of xDbit spatial transcriptome data, we developed a 2-step computational pipeline that integrates raw next generation sequencing (NGS) reads and imaging data (**Fig. 1D**). In the first step of the pipeline (*ReadsToCounts*), spatial coordinates and genetic information were extracted from the raw sequencing reads. After genomic alignment, data were transformed into a spot/gene count matrix. In the second step (*CountsToAnndata*), the SIFT algorithm (Lowe, 2004) was used to register the high-quality and alignment images based on their DAPI channels and calculate a homography matrix. The homography matrix was used to project the xDbit spots onto the high-quality image to generate an integrated AnnData file compatible with Scanpy and Squidpy (Palla et al., 2022; Wolf et al., 2018) for further analysis.

### xDbit performance analysis

To demonstrate the performance improvements of xDbit, we first acquired ST data from murine liver sections using the standard DBiT-seq protocol published by Liu et al. DNA read counts per spot for the liver samples were comparable to the read counts obtained with Dbit-seq on mouse embryo sections (**Fig. 2A**). The lower number of genes per spot for the liver (**Fig. 2B**) sample can be explained by the highly homogeneous cellular composition of the liver, which results in low cell type variance per spot. In the next step, we performed xDbit using two sequentially improved protocols. In the first optimization round, we changed the chemical composition of the initial reactions of the DBiT-seq protocol, namely, the reverse transcription and spatial barcoding reactions. In comparison to DBiT-seq, the reverse transcription reaction, which suffers from low yields (Larsson et al., 2010), was performed on whole tissue sections in the 9-well chip at a concentration of 10 U/μL rather than inside the microfluidic channels to increase the availability of the reverse transcriptase. Furthermore, the concentration of ligase was increased from 15 to 20 U/μL. Spatial barcoding was achieved by two sequential ligation steps, which were performed at lower temperatures and required shorter incubation times than RevT, thus reducing the risk of leakage between channels. Together, the chemistry optimization resulted in a three-fold increase in both read and gene counts per spot compared to DBiT-seq. In the second optimization round, we dehydrated and dried the tissue sections before applying each of the two PDMS chips to improve the attachment of the microfluidic channels. To fill microfluidic channels, equal inlet ports were primed with DNA barcode solutions by centrifugation. Furthermore, bubble traps were added at the transition of the inlets to the microchannels (**Figure S1G**). Collectively, these changes increased the read and gene counts per spot two-fold and four-fold, respectively (**Fig. 2A** and **B**).

**Figure 2.**
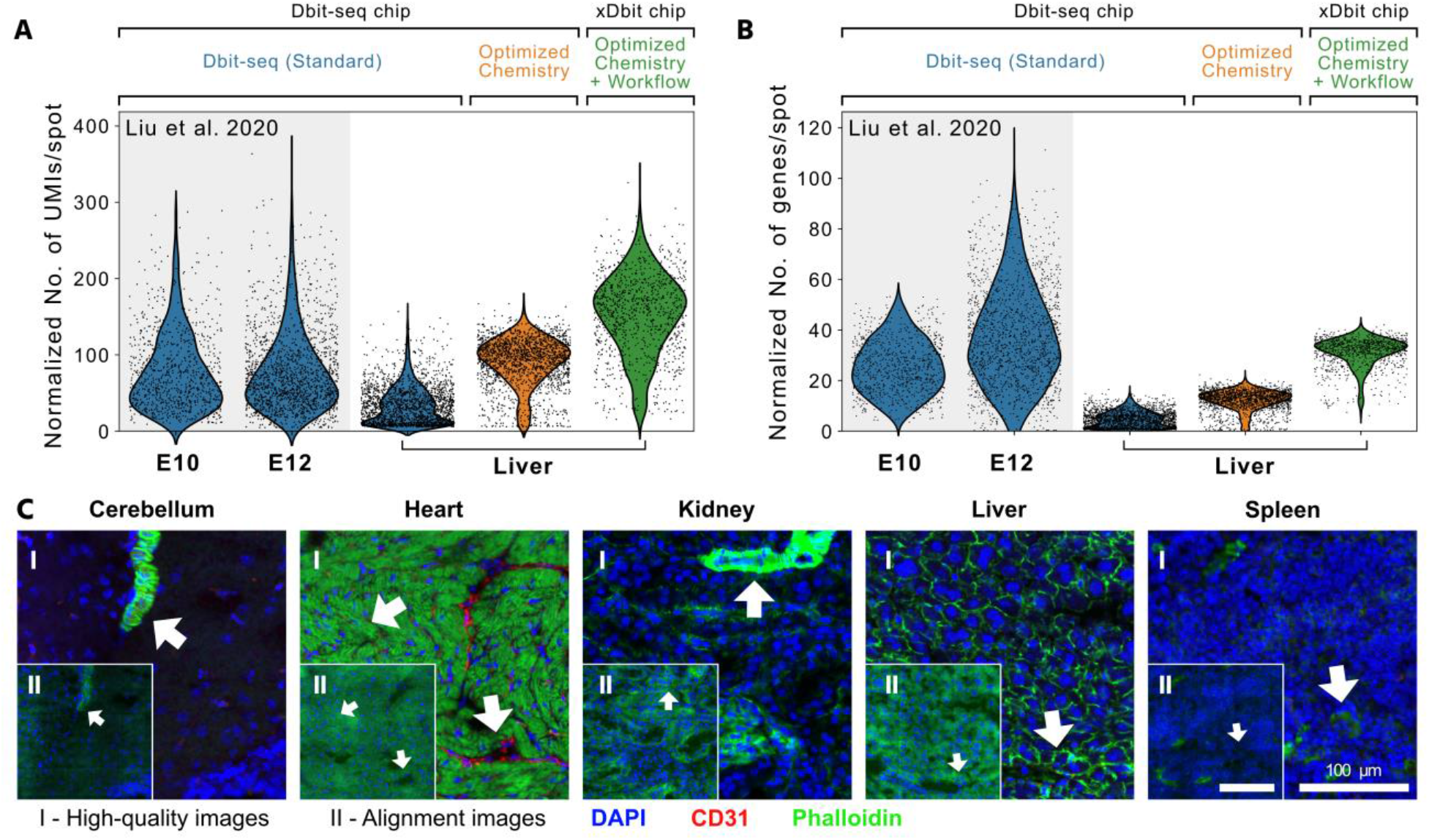
Performance of xDbit and quality assessment of the imaging data. (**A)** and (**B**) Violin plots show the number of UMIs (A) and genes (B) per 50 μm × 50 μm spot, normalized to the sequencing depth (per 10^6^ reads) of the respective experiment. The xDbit method was obtained by sequential optimization of the chemical protocol and workflow of the Dbit-seq method (Standard). For comparison, counts from the previously published Dbit-seq datasets from fresh frozen sections of murine embryos at stage E10 and E12 were added (Liu et al., 2020). (**C**) Representative high-quality (I) and alignment images (II) of mouse tissue sections acquired during xDbit workflow. Blue, red, and green colors denote DAPI, CD31, and phalloidin fluorescence signals, respectively. Scale bar: 100 μm. The quality of the fluorescence signal decreases during the xDbit workflow, which demonstrated the need for the initial high-quality imaging round. UMI: Unique molecular identifier.

It is noteworthy that the structural integrity of the cryo-sections was strongly reduced after the deterministic barcoding workflow because of the physical alignment of the chip platforms to the tissue and enzymatic treatments. Thus, to obtain high-quality image data, which are currently underutilized by standard ST methods (Rao et al., 2021), we acquired imaging data before and after the xDbit workflow. While the images before the xDbit workflow exhibited high-quality features (**Fig. 2C I)**, the images collected after deterministic barcoding steps exhibited the marks required to align the ST data (**Fig. 2C II**). Nuclei integrity was unaffected after the xDbit workflow, and thus alignment images could be registered to high-quality images with the provided *CountsToAnndata* pipeline to transfer the positional information of the alignment marks to the high-quality images (see the Methods section). Taken together, the xDbit workflow provides a multiplexing method for ST and paired high-quality images. Importantly, the xDbit costs for one tissue section are on the order of 125€ (see **Supp. Fig 2H**.).

### Spatially resolved multi-organ dataset with xDbit

To demonstrate the broad applicability of xDbit, we generated high-quality spatially resolved datasets from five different murine organs, including the kidney, heart, cerebellum, spleen, and liver (**Fig. 3**). The UMIs and genes per xDbit spot varied for the organs between 5,000–20,000 and 1,000–5,000, respectively (**Fig. 3A and B**). After dimensionality reduction and transcriptomic clustering using the *Leiden* algorithm (Traag et al., 2019), the projection of xDbit spots onto the respective microscopy images revealed spatially distinct clusters (**Fig. 3C and D**).

**Figure 3.**
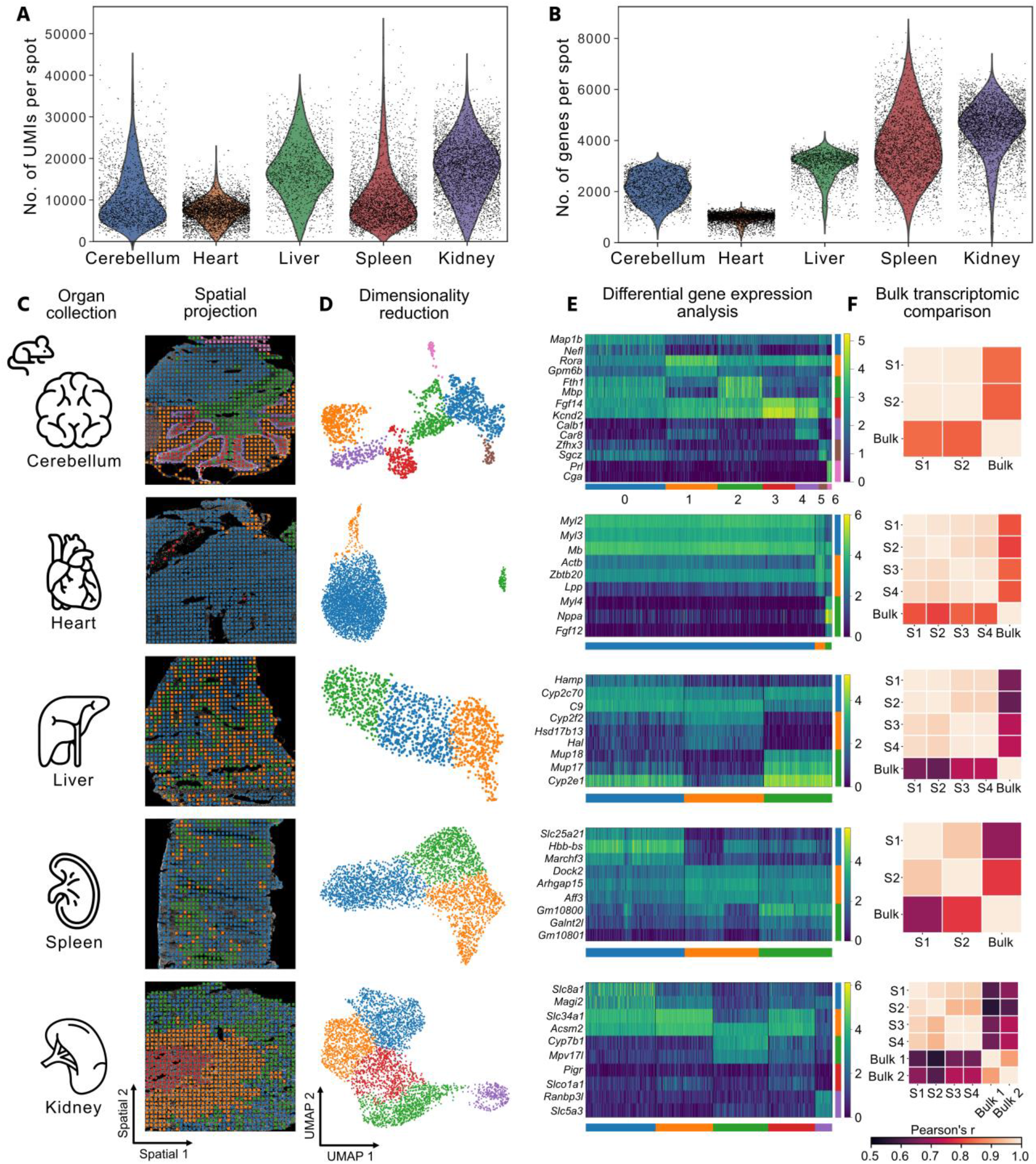
Spatial transcriptomics of multiple murine organs with xDbit. (**A**) and (**B**) UMI and gene counts per spot obtained with the xDbit method for tissue sections of different murine organs. **(C)** Representative DAPI images of five murine organs overlaid with xDbit spatial gene expression. **(D)** Two-dimensional embedding of the xDbit transcriptomic data using UMAP (McInnes et al., 2018). Each spot represents one xDbit spot. Colors denote clusters which were determined using the Leiden algorithm (Traag et al., 2019). **(E)** Top differentially expressed genes of the identified Leiden clusters within the corresponding organs. While the continuous color bar denotes the log-transformed gene expression level, the distinct color bars refer to the Leiden clusters. **(F)** Pearson correlation of tissue bulk transcriptomes from the ENCODE database (Davis et al., 2018; Dunham et al., 2012) with pseudo-bulk xDbit spatial transcriptomic datasets of individual organ samples (S). UMI: Unique molecular identifier; UMAP: Uniform Manifold Approximation and Projection.

Differential gene expression (DGE) analysis between *Leiden* clusters revealed known marker genes for the substructures of the respective organs (**Fig. 3E**). For example, in the heart tissue section, we found the cardiomyocyte markers *Myl2, Myl3*, and *Mb* to be the top differentially expressed genes (Litviňuková et al., 2020; Mantri et al., 2021; Nomura et al., 2018). In the liver section, the zonation markers *Cyp2f2* and *Cyp1a2* were expressed in mutually exclusive areas (Ben-Moshe and Itzkovitz, 2019) indicating that transcriptomic XY resolution is sufficient to define zonated gene expression patterns. In the cerebellar sections of the brain, we were able to identify structures such as the *arbor vitae* (cluster 2) and the cerebellar cortex comprised of clusters 1, 3, and 4 (**Fig. 3 C**). Cluster 4 delineated the course of Purkinje cells in the cerebellar cortex, as confirmed by gene ontology (GO) term enrichment analysis using the STRING algorithm (Franceschini et al., 2013) and the Brenda Tissue Ontology (Gremse et al., 2011) (see **Figure S5A** and **Fig. 3**).

In the spleen, DGE analysis revealed genes that are known to be expressed in the red pulp, such as *Slc25a21* or *Hbb-bs* for cluster 0, and genes expressed in the white pulp, such as *Arhgap15* and *Aff3* for cluster 2. GO term enrichment analysis confirmed the identity of tissue clusters (**Figure S5 D**). To further confirm the high quality of the xDbit datasets, a pseudobulk xDbit dataset was created and compared with published bulk RNA-seq datasets from the ENCODE project (Davis et al., 2018; Dunham et al., 2012). Pearson correlation coefficients between the xDbit pseudobulk and bulk transcriptome data ranged from 0.55–0.83 (**Fig. 3F**).

### Characterization of spatial gene expression

For spatial gene expression pattern analysis of the xDbit ST data, we applied MEFISTO, which is a factor analysis method to identify the driving sources of gene variation in high-dimensional datasets and account for spatial dependencies (Velten et al., 2022). Sections from the same organ showed a comparable number of factors that explained spatial gene expression variations. (**Fig. 4A**). While tissue sections from structurally more complex organs like cerebellum or kidney contained up to six factors explaining the variance in gene expression, in homogenously structured organs like liver or spleen only two factors were sufficient. Investigation of the feature weights of individual factors revealed that the corresponding gene sets influenced the factors in a positive or negative direction (**Fig. 4B**). To further evaluate the performance of MEFISTO on a structurally complex organ, kidney was chosen as model tissue and the first four factors of one kidney section were selected for downstream analysis (dotted frame in **Fig. 4A**). To show that MEFISTO captured structural areas within the tissue sections, we projected the factor values onto the fluorescence image of the respective kidney tissue section (**Fig. 4C**). Factors 1, 2, 3, and 4 define the anatomical regions of the inner and outer medulla, renal tubules in the cortex and medulla, and blood vessels in the kidney, respectively. Similarly, the spatial gene expression of the top positively weighted genes matched the patterns of their corresponding spatial factors (**Figure S6A**). To support factor-to-region assignments, we performed GO term enrichment analysis with the top positively weighted genes of the first four factors (**Fig. 4D**). Analysis of factor 1 showed significant enrichment for terms related to Henle’s loop, a functional structure of the kidney located in the inner medullary region. For Factor 2, the analysis did not show enrichment for specific anatomical regions, but positively weighted genes of this factor were cell type markers for proximal tubules, including *Napsa* and *Serpin1f*. Accordingly, the analysis of positively weighted genes of factor 3 showed significant enrichment in genes of the distal tubules located in the renal cortex. Lastly, the spatial pattern of factor 4 correlated with phalloidin and CD31 staining in the cortical and inner medullary regions of the kidney (**Figure S6 C**). These findings were consistent with the GO term analysis, which showed that genes of the cardiovascular system were enriched. In conclusion, xDbit ST data in combination with MEFISTO factor analysis allowed simultaneous identification and characterization of functional regions in tissue sections from multiple murine organs.

**Figure 4.**
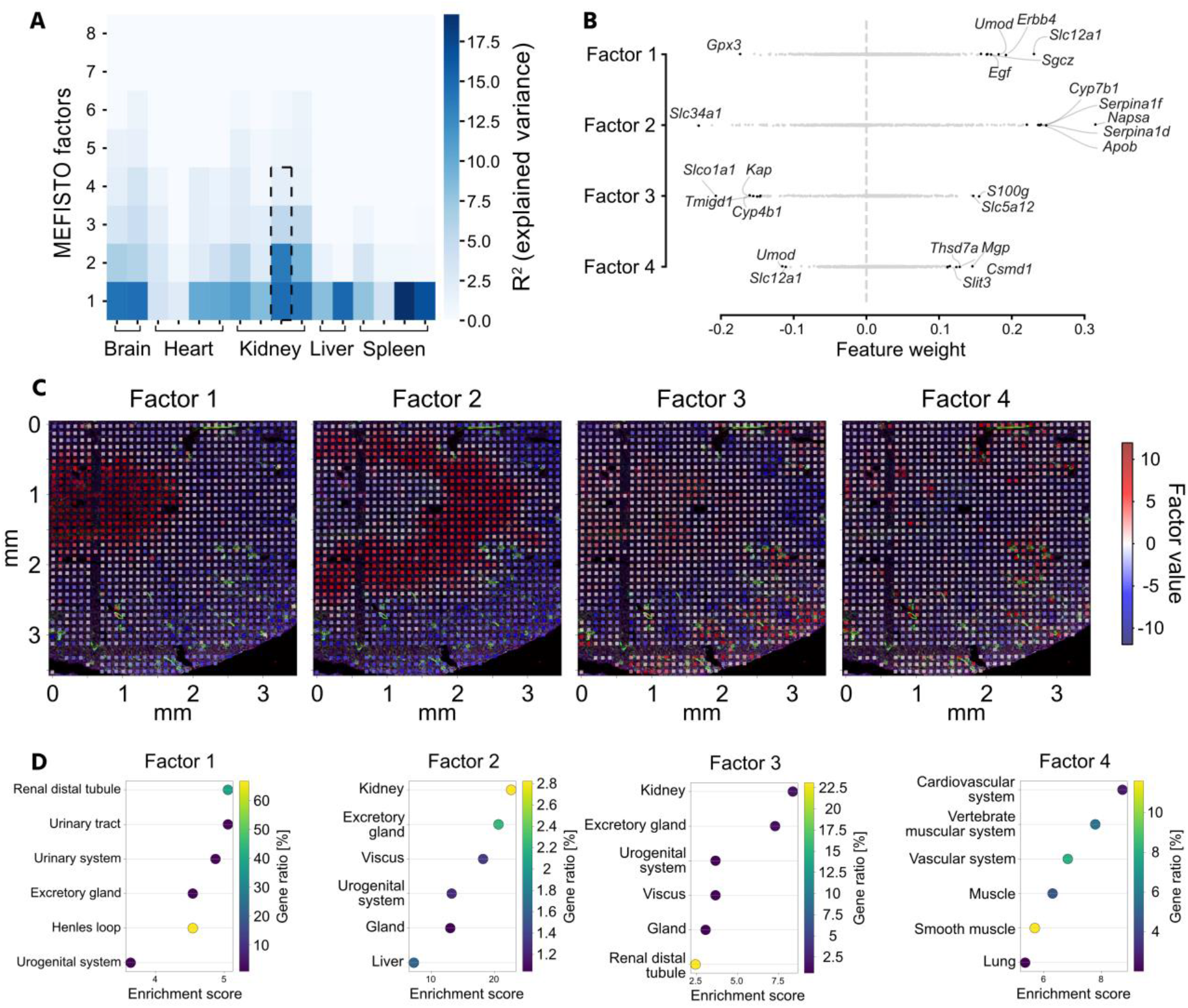
Spatial pattern analysis on xDbit spatial transcriptomic data using MEFISTO factor analysis. (**A**) Results of MEFISTO factor analysis of all individually analyzed xDbit datasets (Velten et al., 2022). Heatmap shows the percentage of explained variance (R^2^) for the first 8 factors of each xDbit replicate. Dotted frame indicates the representative kidney sample which was selected for downstream analysis in **B-D**. (**B**) Scatterplot shows the weight of all genes for the first four MEFISTO factors in the representative sample. Genes with the highest positive or negative weights are labelled. (**C**) High-resolution fluorescence images with overlaid xDbit spots colored for the values of the respective first four MEFISTO factors. Blue: DAPI, Green: Phalloidin; Red: CD31. (**D**) GO term enrichment analysis using the STRING algorithm (Szklarczyk et al., 2019) and the *Brenda Tissue Ontology* database (Gremse et al., 2011). As input we used the top positively weighted genes (> 95% CI) of the first four factors to reveal functional and structural areas of the murine kidney. CI: Confidence interval.

### Deconvolution of xDbit kidney dataset to spatially map cell types

One challenging aspect of ST methodologies and their corresponding computational tools is achieving single-cell resolution across an entire tissue section. For example, existing spatial transcriptomic methods, including *Visium Spatial* (Salmén et al., 2018), *Slide-seqV2* (Stickels et al., 2020), Dbit-seq (Liu et al., 2020), and xDbit, contain multiple cells per spot and are thus unable to reach single-cell resolution. However, single-cell information can be extracted from spatial transcriptomic spots with more than one cell using deconvolution methods (Andersson et al., 2020; Kleshchevnikov et al., 2022; Lopez et al., 2022). In this study, the *cell2location* analysis tool (Kleshchevnikov et al., 2022) was used in conjunction with a published single-cell transcriptome dataset of a murine kidneys (Miao et al., 2021) to obtain the cell-type compositions of each spot on an xDbit kidney ST dataset (**Figure S7A**). The most abundant cell types were cells from the proximal straight tubule (34.6%) and endothelial cells (17.3%), followed by cells from the loop of Henle (15.7%) and the proximal convoluted tubule (10.8%). These findings are in agreement with those of previous studies that investigated the cell type composition of murine kidneys (Chen et al., 2019; Clark et al., 2019). Furthermore, the predicted spatial distribution of these cell types matches the anatomical structure of the kidney (Chen et al., 2019) (**Fig. 5B**). This prediction was further validated by visual correlation of the inferred number of endothelial cells per spot and the fluorescence signal intensity of the endothelial marker CD31 in the kidney section (**Figure S7B**). While cells of the proximal convoluted tubule were found predominantly in the cortex of the kidney, the number of cells of the proximal straight tubule was increased in the outer medulla. Cells of the loop of Henle were mainly predicted to be in the medullar region of the section, which is in agreement with the GO term analysis of MEFISTO factor 1 (see **Fig. 4C and D**). To further challenge the xDbit dataset, we asked whether it is possible to map podocytes, which are cell types located within the glomeruli and have a crucial role in renal filtering processes. High-quality fluorescence images allowed us to identify the position of glomeruli in the tissue section based on phalloidin staining of F-actin, which is a characteristic of glomeruli (Kumaran and Hanukoglu, 2020). The number of inferred podocytes correlated well with the position of the glomeruli, showing high podocyte numbers in spots close to a glomerulus (**Fig. 5 C**). That xDbit spots did not fully align with the glomeruli suggests that the resolution of the spots was larger than the 50 μm × 50 μm area. This might be caused by the diffusion of molecules within the fixed tissue and beneath the covering microfluidic channels. Notably, podocytes are underrepresented in kidney datasets and require special isolation methods (Chung et al., 2020; Karaiskos et al., 2018). The proportion of podocytes detected solely by single-cell transcriptomic data was only 0.3% (Miao et al., 2021) whereas other, less biased studies predicted 3%, a much higher percentage of cells (Clark et al., 2019; Puelles et al., 2016). *Cell2location* inferred a podocyte proportion of 1.7% and thus a more realistic approximation of the kidney cell composition when ST was taken into account (**Fig. 5A** and **Table S8**). In summary, the use of xDbit in conjunction with *cell2location* allows us to map all major renal cell types in a kidney section and generate a more accurate representation of rare cell types in complex microenvironments than scT alone.

**Figure 5.**
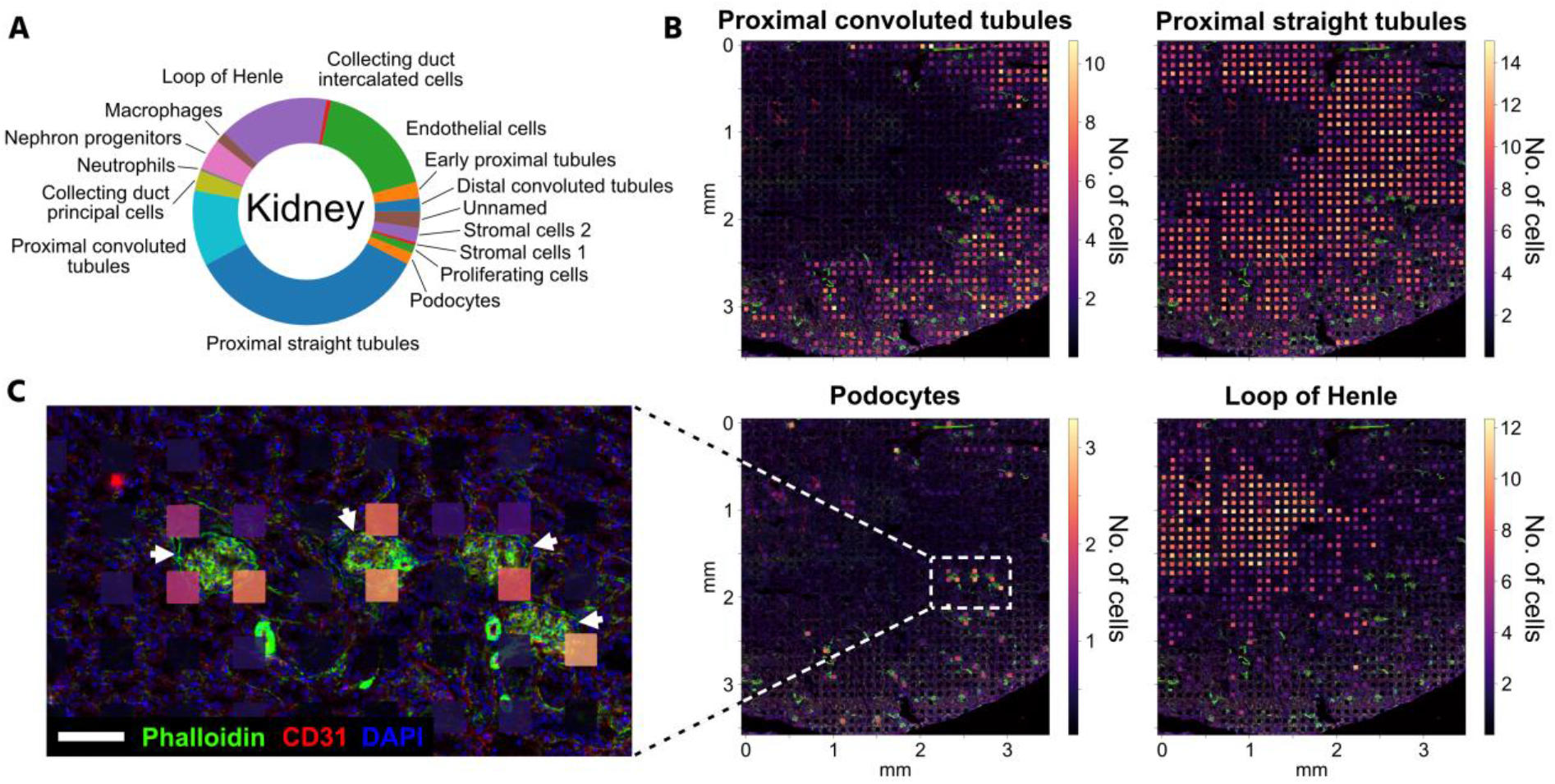
Deconvolution of xDbit kidney dataset to spatially resolve cell types. **(A)** Data of one representative kidney section was deconvolved using *cell2location* (Kleshchevnikov et al., 2022) and a single-cell RNA-seq dataset of the murine kidney (Miao et al., 2021). Pie chart shows the cell type composition of the deconvolved kidney section. **(B)** High-resolution fluorescence images overlaid with xDbit spots colored for the deconvolution results of four representative kidney cell types. Spot colors correspond to the minimum number of cells predicted. **(C)** Detailed fluorescence image. Arrows denote position of glomeruli. Overlaid xDbit spots show the number of podocytes predicted by *cell2location*. DAPI (blue), phalloidin (green), and CD31 (red). Scale bar: 100 μm.

## Discussion

Spatially resolved transcriptomes of tissues from multicellular organisms have greatly expanded our knowledge of complex cellular functions and cell-to-cell communication in healthy and diseased conditions. Single-cell transcriptomics, together with spatial transcriptomics, have become central technologies for mapping cell types in their tissue context and architecture. Most single-cell and spatial transcriptomic studies use a hypothesis-free and explorative design (Bäckdahl et al., 2021; Chen et al., 2020; Mantri et al., 2021; Rao et al., 2021; Wu et al., 2021b). However, to pursue systematic and hypothesis-driven research approaches, ST technologies must comply with the increasing demand of providing multiple replicates per condition, time trajectories, or sampling multiple organs from the same individual at low costs.

In this study, we expanded the technology of deterministic barcoding in tissue to simultaneously analyze nine individual tissue sections. To achieve this, we developed new microfluidic chip platforms to spatially barcode mRNA transcripts in spots with an area of 50 μm × 50 μm. In combination with an optimized chemical workflow, the transcript number of reads and genes per spot were increased by 6 and 12-fold, respectively, compared to the original Dbit-seq method (Liu et al., 2020). Additionally, our results show that xDbit is suitable for different tissues which will facilitate studies focused on complex diseases and multi-organ dysfunction.

Despite the ST technology advances reported here and from others, the lateral diffusion of molecules in the barcoding step of ST methods, limits the resolution of sequencing-based ST methods to the range of 5-10 μm (Chen et al., 2022). Furthermore, ST datasets with subcellular resolution require elaborate algorithms to segment single cells based on the spatial transcriptome (Chen et al., 2015). Rather than further increasing the resolution of spatial transcriptomic methods, an alternative approach is the use of single-cell transcriptomic datasets and computational methodologies to increase the resolution of the datasets *in silico*. Thus, a large and complex experimental design with the objective of mapping cell transcriptomes and retaining tissue context requires more affordable technologies. Here, we have shown that xDbit is a low-cost ST technology (ca. 125 € per sample) that provides robust and accurate analysis of spatial gene expression patterns. The achieved transcript read depth on xDbit spots together with deconvolution tools is sufficient to resolve rare cell types, such as podocytes in the glomeruli of the mural kidney. Thus, xDbit is an ST methodology that optimally balances the cost and throughput. Further engineering efforts will focus on increasing the screening areas, in addition to read depth. The xDbit workflow could be further scaled to larger screening areas by increasing the microfluidic channel length as well as the microfluidic chip platform. The only limiting factor of the xDbit approach is the fluid resistance, which scales linearly with channel length. A higher fluid resistance requires a higher fluid forward pressure to drive fluid flow, which in turn can induce leakage between the microchannels and disruption of the underlying tissue.

Furthermore, barcoding strategies with microfluidic channels can be combined with a multitude of modalities, including DNA-barcoded antibodies (Liu et al., 2022), chromatin accessibility (Deng et al., 2022b), and epigenomic readouts (Deng et al., 2022a). To increase adaptability, xDbit libraries can be sequenced using standard next generation sequencing platforms.

Since the microfluidic workflow has adverse effects on the integrity of the tissue sections and image information is needed to further enhance the power of spatial transcriptomic data (Rao et al., 2021), we introduced two imaging steps to allow the acquisition of high-quality imaging data. This allows the platform being used for the analysis of high-resolution image features in conjunction with transcriptomic information. In addition to technical advances, we have provided an open-source analysis pipeline to generate xDbit datasets and make the method easily accessible. This includes a semi-automatic image registration pipeline and the introduction of alignment markers to robustly align the fluorescent images with ST data. In summary, using xDbit, we expanded the toolbox of spatial transcriptomic methods for higher throughput measurements and improved both the transcriptomic and imaging quality of the resulting datasets.

## Methods

### Ethics statement

Animal experiments were carried out in compliance with the German Animal Protection Act and with the approved guidelines of the Society of Laboratory Animals (GV-SOLAS). All animals used within this study were kept at the Helmholtz Zentrum AVM Research Unit Comparative Medicine, Neuherberg, Germany. This study and the animal facilities were approved by the Institutional Animal Welfare Officer and by the Government of Upper Bavaria, Germany.

### Husbandry and tissue collection

Wild-type C57BL/6J mice were purchased from Charles River UK Ltd (Margate, United Kingdom) and were maintained under specific pathogen-free conditions under strict 12 h dark-light cycles. All mice were kept in a positive pressure system, maintaining a temperature between 19 and 23°C, 55% humidity, and had free access to water and standard mouse chow diet.

Three male C57BL/6J mice (age 3-4 months) were used in this study. At the time of experiment, mice were sacrificed in accordance to GV-SOLAS regulation, and were subsequently dissected. Heart, liver, kidney and spleen were collected from the same two mice while the brain sample was collected from a different mouse. The organs cryo preserved using Tissue-Tek OCT Compound (CellPath Ltd, UK) into Tissue-Tek Cryomolds (Sakura Finetek, USA). All cryo embeddings were frozen in pre-chilled 2-methylbutane on dry ice. After freezing, cryo embeddings were transferred into −80°C freezer for long term storage. For the brain, cerebrum and cerebellum were embedded separately.

### Master mold fabrication

Master molds for the horizontal and vertical xDbit chips were fabricated according to standard SU-8 (SU-8 3050; Microresist Technology, Germany) photolithography protocols.(Unger et al., 2000) To prevent PDMS adhesion to the SU-8 mold, the surface was spin-coated with a thin film (<1 μm) of CYTOP^™^ (AGC Chemicals, Japan).To evaporate the CYTOP^™^ solvent the SU-8 mold was heated to 160 °C for 1 h.

### xDbit chip fabrication

Horizontal and vertical xDbit chips were manufactured by casting a 5 mm PDMS (Sylgard® 184, Dow Corning, MI, USA) layer (ratio 10:1 of base material to curing agent) onto the SU-8 master mold. After degassing for 1 hour in an evacuated desiccator at room temperature (rt), the PDMS was cured for 1 hour at 80 °C. The cured PDMS chip was peeled off, cut into the required size and inlets and outlets were punched using a 14 gauge needle.

### Fabrication of non-PDMS modules

To press the xDbit chip onto the tissue sections, a plastic clamp was milled in acrylic glass. Well modules that allow the precise application of reagents onto the tissue sections and molds for PDMS gaskets were 3D-printed with a DLP stereolithography printer (Pico2HD, Asiga, Australia) using the resin PlasGRAY (Asiga, Australia). Printing parameters including light intensity and exposure time were set according to the manufacturer’s material file. After exposure, the printed part was removed and sonicated in isopropanol for 10 minutes. Afterwards, the printed parts were incubated for 4 hours at room temperature to remove excess isopropanol and post-cured at 2000 flashes per side (Otoflash curing unit).

### Fabrication of PDMS modules and gaskets

PDMS gaskets and the vacuum adapter were manufactured by replica molding using 3D printed molds. After 3D printing as described above, the molds were dip-coated with CYTOP^™^ (AGC Chemicals) and incubated on a hotplate for 8 h at 80 °C. For gaskets, a 5 mm layer of PDMS (ratio 10:1 of base material to curing agent) was poured into one well of a 6-well plate. The mold was pressed upside down into the PDMS and the material was degassed for 1 hour in an evacuated desiccator at rt. After curing for 1 hour at 80 °C, the PDMS gasket was peeled off carefully and excess material was removed with a knife. For the vacuum adapter, the mold was glued to the bottom of a well and PDMS was poured over the mold. Degassing and curing was performed as described and a hole was punched in one of the sides using a 2 mm punching needle. An overview of all modules needed in the xDbit workflow is shown in **Figure S1 A-C**.

### Tissue preparation

For sectioning, the organs were warmed to −15—18 °C inside the cryostat (Leica). Object slides with marked placement areas were cooled inside the cryostat before use for at least 5 minutes. The tissue blocks were sectioned with a thickness of 10 μm using RNAse free equipment, placed in predetermined positions on the object slide (**Figure S1H**) and attached by warming the backside of the object slide with a finger. The sectioned samples were stored at −80 °C.

### Generation of optimal RevT primer barcode sets

To prevent reverse transcription bias from RevT primer barcodes, we used mixes of multiple RevT primers in the RevT reaction. BARCOSEL (Somervuo et al., 2018) was used to generated 9 sets of RevT primers with 4 barcodes per set (**Table S2**). In this study, we did not barcode the wells in the RevT step separately and instead mixed sets 1-4 to further increase the diversity. The RevT primers (**Table S1**, Sigma) were dissolved in ultrapure water at a concentration of 100 μM and mixed at this concentration.

### *Preparation of* Ligation Barcoding Plates

Barcoded ligation oligos (**Table S3 and**Fehler! Verweisquelle konnte nicht gefunden werden., Sigma) were dissolved in ultrapure water at a concentration of 100 μM and stock plates were stored at −20 °C. Separately for ligation round #1 and ligation round #2, 36 ligation barcode oligos were annealed with the respective bridge oligo as described in **Table S5**. In brief, 21.1 μL of a bridge oligo (1 mM, Sigma) were mixed with 296 μL water and 317 μL 2X annealing buffer (5 mM Tris, 100 mM NaCl) to a final concentration of 33.33 μM. Then, 4 μL of one ligation barcode oligo (100 μM) and 12 μL of the diluted linker were mixed in a 96 well plate. Using a PCR cycler, the oligos were denaturated at 95 °C for 2 minutes and cooled to 20 °C at a rate of −0.1 °C/s to anneal the strands. The annealed oligo stock plates were stored at 4 °C for short-term or −20 °C for long-term storage. Before the experiment, 1 μL of each barcode was distributed to fresh PCR plates, later called *Ligation Barcoding Plate* #1/#2.

### Fixation, permeabilization and blocking

The object slide with tissue sections was thawed at 37 °C for 1 minute on a heated plate. Clamp, 1-well top module and PDMS gasket (**Figure S1E**) were assembled, aligned and attached to the tissue slide. The tissue sections were washed with 1X RNAse-free phosphate buffered saline (PBS, Invitrogen) supplemented with Murine RNAse inhibitor (1 U/μL, “RI”, New England Biolabs) and ribonucleoside vanadyl complex (RVC, 10 mM, New England Biolabs) and fixed in 4% paraformaldehyde (PFA, Sigma) for 40 minutes at room temperature (RT). After three washes in 1X PBS complemented with RVC (10 mM, “PBS+RVC”), the tissue sections were permeabilized with 0.2% Triton X-100 (Sigma) in PBS+RI for 10 minutes at RT and blocked for 30 minutes at RT with 1% bovine serum albumin (BSA, Thermo Fisher).

### Staining and high-resolution confocal imaging

The CD31 primary antibody (Thermo Fisher, PA5-16301) was diluted 1:50 in antibody diluent (PBS+RI supplemented with 0.1% Tween-20 and 3% donkey serum), added to the sections and incubated for 30 min at RT. After 3X wash in PBS-T (0.1% Tween-20) supplemented with RVC (PBS-T+RVC), nuclei, actin filaments and primary antibody were stained using DAPI (1.25 μg/mL, Sigma), Phalloidin-iFluor647 (1.25X, Abcam) and AF555 secondary antibody (Invitrogen, A-31572) in antibody diluent for 30 minutes at room temperature in the dark. The tissue sections were washed three times in PBS-T supplemented with RI (PBS-T+RI) and mounted in 85% ultrapure glycerol (Sigma) supplemented with 2 U/μL RI using #1.5 coverslips (Menzel). Images were acquired using an LSM 880 confocal microscope (Zeiss) and a 20X/0.8 objective (Zeiss) at a final resolution of 0.24 μm/pixel.

### Reverse transcription (RevT)

The coverslip was removed by holding the object slide in a 45° angle with the coverslip facing down into 3X saline sodium citrate (SSC) buffer until the coverslip falls off. The sections were dipped 3X in ultrapure water and dried under airflow. Clamp and 9-well module (**Figure S1F**) were assembled, aligned and attached to the tissue slide. PBS+RI supplemented with 1% BSA was added and stored at 4 °C for maximum 30 minutes until the next steps were performed. An RevT reaction mix was prepared from 514.8 μL ultrapure water (Thermo Fisher), 158.4 μL RevT buffer (5X, Maxima H Minus RT Kit, Thermo Fisher), 39.6 μL dNTPs (10 mM, New England Biolabs), 19.8 μL RI, 19.8 μL RevT primer set and 39.6 μL Maxima H Minus Reverse Transcriptase (200 U/μL, Thermo Fisher). 80 μL of the mix were added to each well, the wells were sealed and the slide was incubated for 30 minutes at RT and 90 minutes at 42 °C in a closed thermoshaker without agitation. To ensure equal heat distribution and minimize evaporation an aluminum block was placed between object slide and hot plate and wet tissues were added to the closed container. Afterwards, the tissue sections were washed once in PBS-T+RVC and the 9-well module was removed.

### Spatial barcoding by ligation (horizontally or vertically)

The object slide was dipped 3X into ultrapure water to remove salts and the tissue sections were dehydrated stepwise by incubation in 70, 85 and 99.5% ethanol for 1 minute each and dried briefly under airflow. The horizontal (ligation round #1) or vertical (ligation round #2) PDMS xDbit chip was aligned, attached to the tissue sections and placed into the clamp (**Figure S1D**) and the screws were tightened uniformly and strongly to prevent leakage.

To rehydrate the tissue, 5 μL of PBS+RI were added to each inlet and the channels were filled by applying 300 mbar vacuum to the outlets using a PDMS vacuum adapter (**Figure S1G**) and incubated for about 10 min at RT. A ligation reaction master mix was prepared from 149.66 μL ultrapure water, 26.3 μL T4 DNA Ligase buffer (New England Biolabs), 2.51 μL 10% Triton X-100 (Sigma), 13.1 μL Murine RNAse inhibitor, 5.25 μL Tartrazine (10 mg/mL, Carl Roth) and 13.2 μL T4 DNA ligase (New England Biolabs). 4 μL of the master mix were added to the *Ligation Barcoding Plate* #1 or #2 (see above) respectively for a total of 5 μL and centrifuged down briefly.

The inlets of the xDbit chip were emptied using a vacuum aspirator with attached pipette tip and 5 μL of each barcode was added to the inlets according to the inlet filling scheme (**Table S6**). The outermost channels were filled with an alignment marker mix consisting of 80 μg/mL anti-BSA antibody (Invitrogen) in antibody diluent. To remove air bubbles in the inlets, the chip was centrifuged at 100 x g for 1 min. The channels were filled using vacuum as described before. Inlets and outlets were sealed and the chip was incubated at 37 °C in a closed thermoshaker without agitation. To ensure equal heat distribution and minimize evaporation an aluminum block was placed between object slide and hot plate and wet tissues were added to the closed container. After 15 min the vacuum was applied again to remove air bubbles in the channels and the chip was incubated another 15 min at 37 °C for a total of 30 min reaction time. The inlets were emptied with the vacuum aspirator and the channels were washed for 5 min with PBS-T+RI. Afterwards, the channels were emptied and the chip was removed.

### Secondary staining and alignment imaging

The alignment markers were stained with 4 μg/mL donkey anti-rabbit AlexaFluor 555 secondary antibody (Invitrogen, A-31572) in PBS-T+RI supplemented with 3% donkey serum, 1.25 μg/mL DAPI and 1.25X Phalloidin-iFluor647 for 30 minutes at room temperature in the dark. Afterwards, the tissue sections were washed three times in PBS-T+RI and mounted as described before. Images were acquired using an LSM 880 confocal microscope (Zeiss) and a 20X/0.8 objective (Zeiss) at a final resolution of 0.49 μm/pixel using the fastest possible scanning mode.

### Lysis and sample collection

The coverslip was removed from the tissues and the 9-well module attached as described before. Lysis buffer was prepared from 10 mM Tris-Cl pH 8.0, 200 mM NaCl (Sigma), 50 mM EDTA pH 8.0 (Life Technologies), 2% SDS (Bio-Rad) and 2 mg/mL proteinase K (New England Biolabs). The tissue sections were lysed separately in 75 μL lysis buffer for 2 hours at 55 °C. To prevent evaporation, the wells were closed with a PDMS piece which was fixed with tape and incubation was conducted in a closed container containing wet tissues. Afterwards, possibly remaining parts of the tissue sections were scraped off with the pipette tip and the lysates were collected in nine separate DNA LoBind tubes (Eppendorf). The wells were washed once with 40 μL of lysis buffer and the washing solution was pooled with the lysate. Samples were stored at −80 °C.

### cDNA purification

396 μL of Dynabeads MyOne Streptavidin C1 (44 μL per sample, Thermo Fisher) were washed three times in 800 μL 1X B&W buffer (see manufacturer’s manual) supplemented with 0.5% Tween-20 and 0.05 U/μL RI and resuspended in 950 μL of 2X B&W buffer supplemented with RI (100 μL + 5% per sample). The lysates were thawed at rt, brought to 100 μL with ultrapure water and 5 μL PMSF (200 mM, Cell Signaling) were added and incubated for 10 minutes at rt to block Proteinase K activity. To bind the cDNA to the beads, 100 μL of the resuspended Dynabeads were added to the lysates, vortexed and incubated for 1 hour at rt under agitation (1200 rpm). Afterwards, the beads were washed two times in 1X B&W-T + RI for 5 min at rt under agitation. Likewise, a final washing step was performed in 10 mM Tris-Cl pH 8.0 buffer supplemented with 0.01% Tween-20.

### Template switch

A template switching reaction mix (TSR mix) was prepared from 360 μL ultrapure water, 180 μL RevT buffer (5X), 180 μL Ficoll PM-400 (20%, Sigma), 90 μL dNTPs (10 mM), 22.5 μL Murine RNAse inhibitor, 22.5 Template Switching Oligo (“TSO”, 100 μM, Ella Bioscience) and 45 μL Maxima H Minus Reverse Transcriptase (200 U/μL). The beads with the bound cDNA were placed against a magnetic rack and washed once in ultrapure water. Then, the beads were resuspended in the TSR mix and incubated for 30 min at RT and 90 minutes at 42 °C under agitation (1200 rpm). Afterwards, the samples were placed against a magnetic rack and washed once in ultrapure water.

### PCR amplification

A PCR mix was prepared from 869 μL ultrapure water, 1034.6 μL Kapa Hifi 2X Master Mix (Roche), 82.8 μL cDNA amplification forward primer (10 μM, oSR321211_TSO_fwd) and 82.8 μL reverse primer (10 μM, oSR321212_TSO_rev, see **Table S5**). Each sample was resuspended in 220 μL PCR mix and split equally into 4 different PCR tubes. PCR was performed using following program: 95 °C for 3 min, then 5 cycles of 98 °C for 20 s, 65 °C for 45 s and 72 °C for 3 min. Afterwards, the reaction mixtures were pooled and placed against a magnetic rack. 200 μL of each sample were transferred to a fresh tube and 2 μL of SYBR Green qPCR dye (100 μM, Jena-Bioscience) were added. To account for differences in the cDNA content between the samples an optimal number of PCR cycles was determined for each sample separately. Duplicates of 10 μL of each sample were transferred into a qPCR plate and measured in a Viia 7 qPCR machine (Applied Biosystems) using following program: 95 °C for 3 min, then 40 cycles of 98 °C for 20 s, 67 °C for 20 s, 72 °C for 1 min. The optimal cycle number was defined as the cycle where the qPCR curve reaches 25% of its maximum intensity. The remaining 180 μL per sample were distributed into two PCR tubes and the following qPCR program was run with the previously determined cycle number n: 95 °C for 3 min, then n cycles of 98 °C for 20 s, 67 °C for 20 s, 72 °C for 3 min, and a final extension at 72 °C for 5 min, then hold at 4 °C. Afterwards, the qPCR reactions were pooled per sample.

### cDNA purification

The amplified cDNA was purified using SPRIselect^™^ beads (Beckman Coulter) following a left sided size selection with a bead-to-sample ratio of 0.8X. In brief, 160 μL of sample were mixed with 128 μL of resuspended SPRIselect beads and incubated for 5 min at rt. Beads were washed two times in 85% ethanol and air-dried for 3 min. The cDNA was eluted in 20 μL ultrapure water by incubation at 37 °C for 10 min. The supernatants were transferred to a fresh tube resulting in 9 tubes of purified cDNA. The quality of the cDNA was analyzed using the Bioanalyzer High Sensitivity DNA chip (Agilent) and samples were stored at −20 °C.

### Library preparation and sequencing

The concentration of the cDNA was determined using a Qubit 1X dsDNA assay (Invitrogen) and the sequencing library was generated using the Nextera XT DNA Library Preparation Kit (Illumina). The quality of the library was assessed using the Bioanalyzer High Sensitivity DNA chip (Agilent). Samples were sequenced on a NovaSeq 6000 system (Illumina) at a sequencing depth of minimum 50,000 reads per spot using a 100 cycles kit in paired-end mode. Following read length configurations were used: R1: 7 cycles, i7: 6 cycles, R2: 58 cycles.

### Pipeline overview

Integration of sequencing results and imaging data was performed using a custom pipeline which is published open-source on Github (https://github.com/jwrth/xDbit_toolbox) and combines two previously published analysis pipelines: Drop-seq tools v2.1.0 (Nemesh, 2018) and splitseq_pipeline (Wegmann, 2019) with custom Python and bash scripts scripts. The pipeline consists of 2 main steps: (1) *ReadsToCounts* and (2) *CountsToAnndata* (**Fig. 1D**). The first part of the pipeline needs to be run on a Linux machine while the second part was tested both on a Linux and Windows machine. In the following sections the pipeline is explained briefly. Detailed instructions to process xDbit data can be found in the Github repository. For plotting the Python packages *matplotlib* v3.5.1 (Hunter, 2007) and *seaborn* v0.11.2 (Waskom, 2021) were used. Image transformations were predominantly performed using the *OpenCV* package (Bradski, 2000).

### ReadsToCounts

This script takes two FASTQ files (Read 1 and Read 2) and barcode-coordinate information as input and processes them as follows: Read 1 sequences are trimmed and filtered using *cutadapt* v3.7 (Martin, 2011) and mapped against the mm10 (GRCm38) mouse genome using STAR-2.7.4a (Dobin et al., 2013). Unique molecular identifiers (UMIs) and spatial barcodes are extracted from Read 2 using the Drop-seq tool *TagBamWithReadSequenceExtended* and a custom Python 3 pipeline assigns coordinates using barcode information provided in a CSV file. The *DigitalExpression* function is used to collapse the UMIs and generate a spot-count matrix. RNA metrics are calculated using *CollectRnaSeqMetrics*.

### CountsToAnndata

In this step the spot-count matrix and imaging data are aligned and integrated. In brief, the positions of the alignment marker vertices are extracted semi-automatically from the alignment images using *napari* (Sofroniew et al., 2022) and *Squidpy* (Palla et al., 2022). The coordinates of the vertices are used to register alignment image and xDbit spots by performing an affine transformation using *OpenCV* (Bradski, 2000). In order to align the high-resolution images of the first imaging round with the xDbit spots, the SIFT algorithm (Lowe, 2004) is used to extract common features between the alignment DAPI image and the high-resolution DAPI image. Based on the coordinates of these features an affine transformation matrix is determined, which is used to align the xDbit spots to the high-resolution image. The dataset is saved in the *AnnData* format (Virshup et al., 2021). In this study we included intronic reads (**Figure S4**) into the analysis.

### Preprocessing

Pre-processing of the count matrices was performed using the Python 3 tools Scanpy v1.8.2 (Wolf et al., 2018) and Squidpy v1.1.2 (Palla et al., 2022). To remove the background, we excluded spots with DAPI signal below a threshold. The removed background spots were used to filter out all genes that had a mean background expression, μ_b_, below a threshold, t_g_, defined as:

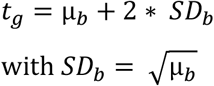

We assumed a Poisson distribution of the background read counts and calculated an approximation of its standard deviation SD. All further analyses were performed according to current best practices in single-cell RNA-seq and Spatial Transcriptomics analysis (Luecken and Theis, 2019; Palla et al., 2022) and can be reproduced using Jupyter Notebook (for more information see Data and Code availability). Counts were normalized, log-transformed, and the top 2000 highly variable genes were determined.

### Dimensionality reduction and clustering

For visualization in lower dimensional space, we calculated the top 50 principal components and generated a two-dimensional representation using Uniform Manifold Approximation and Projection (UMAP) (McInnes et al., 2018). To group the spots into transcriptomically similar clusters the Leiden algorithm (Traag et al., 2019) was applied. Overlay plots of transcriptomic spots and image data were generated using a custom plotting function.

### Differential gene expression analysis

Differentially expressed genes for each leiden cluster were calculated by applying Scanpy’s *rank_genes_groups* using the Wilcoxon rank-sum test and default settings. The top 3 differentially expressed genes were visualized using *rank_genes_groups_heatmap*. For downstream analyses the 300 most significantly differentially expressed genes were used. Information about protein expression of differentially expressed genes in the respective tissues has been taken from The Human Protein Atlas (Mathias et al., 2015).

### Gene Ontology (GO) term enrichment analysis

For GO term enrichment analysis we used APIs of the STRING web server (Szklarczyk et al., 2019). A detailed description on the how the enrichment is calculated can be found in (von Mering et al., 2005). The resulting False Discovery Rate (FDR) shows p-values corrected for multiple testing using the Benjamini-Hochberg procedure. Enrichment scores are represented as negative log_10_ of the FDR. For our analysis we searched for enrichments in the *Brenda Tissue Ontology* database (BTO) (Gremse et al., 2011) and the *Biological Processes* GO database (Ashburner et al., 2000; The Gene Ontology Consortium, 2021).

### Correlation with bulk sequencing data

To compare xDbit ST data with published bulk sequencing data we generated a pseudobulk dataset of the xDbit dataset by summing up the counts of all spots per gene. Following bulk sequencing datasets were downloaded from the ENCODE project website (Davis et al., 2018; Dunham et al., 2012): ENCSR000CGZ (Heart); ENCSR000CHA (Kidney); ENCSR000CGW and ENCSR966JPL (Spleen); ENCSR000CHB (Liver); ENCSR000CGX (Cerebellum). Both the bulk and the pseudobulk datasets were normalized to transcripts per million (TPMs) and log transformed. To analyze the correlation of datasets per organ the Pearson correlation coefficient was calculated pairwise and results were visualized as heatmap.

### Comparison with published Dbit-seq datasets

Previously published Dbit-seq datasets from embryonic sections (Liu et al., 2020) were downloaded from the Gene Expression Omnibus database with the accession code <u>GSE137986</u>. Of the whole dataset following experiments were retrieved for the comparison: “GSM4189613_0702cL” (Embryo stage E10 – 162,684,631 raw reads) and “GSM4189612_0628cL” (Embryo stage E12 – 53,619,846 raw reads). In addition to the xDbit datasets, we used for comparison (1) a dataset that was generated in-house following the protocol of the original Dbit-seq method and (2) a dataset that was generated using the original Dbit-seq chip without serpentine channels but with the optimized biochemical protocol of xDbit. All datasets were normalized to the total number of raw sequencing reads and then compared by the normalized values of total counts per spot and number of genes per spot.

### Image processing

For image processing and generation of figures, we used Fiji ImageJ v1.53c (Schindelin et al., 2012) and the Quickfigures toolkit (Mazo, 2021). Stitching of the tiled images was performed using a custom ImageJ script utilizing the Grid/Collection Stitching algorithm (Preibisch et al., 2009).

### MEFISTO factor analysis

MEFISTO factor analysis (Velten et al., 2022) was performed using the Python package *mofapy2* (v0.6.4). Spatial spot coordinates were used as covariates and only highly variable genes were selected for the analysis. MEFISTO was run on each tissue section separately using following parameters: factors=10; frac_inducing: 0.5; sparseGP=True; start_opt=10; opt_freq=10. Models were saved as hdf5 files and downstream analysis was performed using the *mofax* toolbox (https://github.com/bioFAM/mofax). To investigate the first four factors functionally, for each factor the top weighted genes (> 95 confidence interval) were selected and used for GO term enrichment analysis using the STRING algorithm (Szklarczyk et al., 2019) as explained above.

### Cell type mapping in xDbit kidney data

To map the cell types from single-cell datasets onto xDbit spatial transcriptomics data of the murine kidney, we applied *cell2location* (v0.1, Kleshchevnikov et al., 2022). The single-cell RNA-seq dataset was retrieved from a previous publication including P0 and adult mice samples (Miao et al., 2021, GEO: GSE157079). For the analysis, only cells from adult mice were selected and mitochondrial genes were removed from both the single-cell and the representative xDbit kidney dataset. Genes were filtered using the cell2location gene_filter function, filtering out genes that were detected in less than five cells and less than 0.05 % of cells. Anndatas were prepared for analysis using scvi-tools (v0.16.4, Gayoso et al., 2022). The single-cell model to infer expression signatures of cell types was trained in 250 epochs. Spatial mapping was performed with default parameters, except for: N_cells_per_location=20; detection_alpha=20; max_epochs=30000; batch_size=None; and train_size=1. To show the minimum number of cells, we used the 5 % quantile of the resulting posterior distribution, reflecting the confidently predicted number of cells.

## Supporting information

Supplementary Tables S1-S4 Oligos

## Data and code availability

All raw sequencing data and preprocessed xDbit data including spatial transcriptomic data with aligned images are available on GEO under the accession number GSE207843.

All other data including notebooks, functions and environment files with package versions to rerun the analysis are available in the Github repository https://github.com/jwrth/xDbit_toolbox. The computational pipeline, consisting of the scripts *ReadsToCounts* and *CountsToAnndata* can be found in the subfolders with the respective names. ImageJ scripts to stitch images from tile scans are deposited in the folder named ‘imagej’. CAD plans to manufacture the xDbit master molds using photolithography as well as the plans of the 3D-printed and milled parts necessary for the workflow are stored in the ‘cad’ folder.

## Acknowledgements

We thank Thomas Gerlach and Josef Promoli from the workshop at Helmholtz Zentrum München for milling acrylic glass parts. This work was supported by the Helmholtz Pioneer Campus and ERC Consolidator Grant (Number 772646). We thank Inti I. A. de la Rosa Velazquez and G. Eckstein for performing the NovaSeq sequencing at the Bioinformatics Core Facility of the Helmholtz Zentrum München. We thank T. Walzthöni for the bioinformatics support. We thank S. Kublik for performing the NextSeq sequencing at the Research Unit Comparative Microbiome Analysis of the Helmholtz Zentrum München.

## Author contributions

J.W., N.C., C.M., and M.M. designed the study. J.W. and N.C. designed the xDbit microfluidic chip platform and the 3D-printed modules. N.C. and S.B. manufactured the microfluidic chip and the 3D-printed modules. J.W. and S.B. developed the experimental method of xDbit. J.W. developed the imaging strategy and the computational pipeline. K.Y. and S.C. collected the murine organ samples. J.W. performed all xDbit experiments, processed the raw data and performed all downstream computational analyses. M.M. and C.M. received the funding and supervised the study. The manuscript was written by J.W. and M.M. All authors corrected and approved the manuscript.

## Supplemental information

### Supplementary Figures

**Figure S1.**
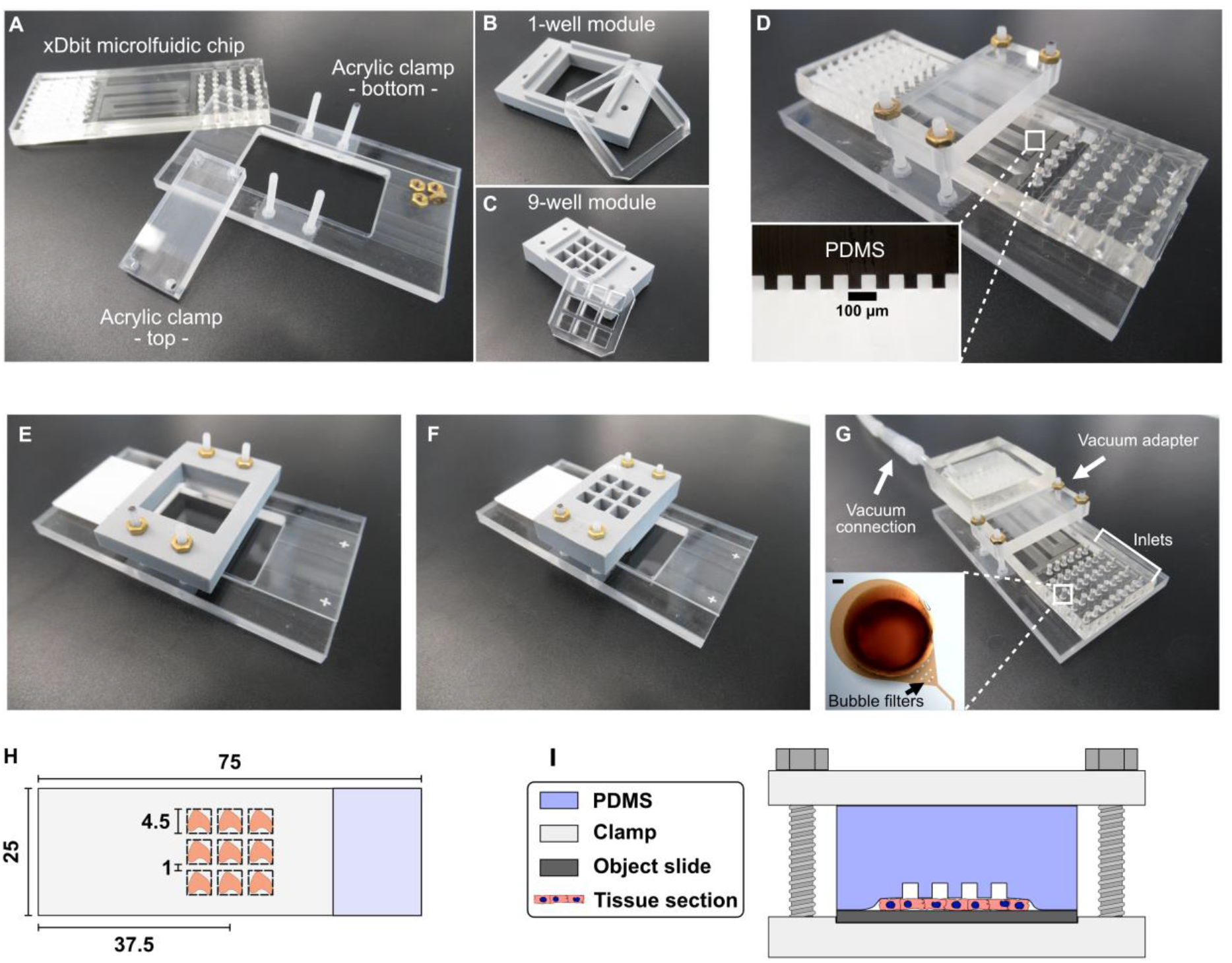
Overview of xDbit components. **(A)** All components required to perform spatial barcoding using the xDbit microfluidic chip. **(B)** Image of 3D-printed 1-well module and respective PDMS gasket to treat all tissue sections at once. **(C)** Image of 3D-printed 9-well module and respective PDMS gasket used to perform the reverse transcriptions and lysis steps. **(D)** xDbit microfluidic chip with assembled clamp. Detail image shows cross section of microfluidic channels with heights and widths of 50 μm. Scale bar: 100 μm. **(E and F)** Assembled 1-well and 9-well modules with clamp, respectively. **(G)** Assembled xDbit microfluidic chip, clamps, and attached vacuum adapter. Enlarged image shows one inlet filled with food dye-colored water to illustrate the bubble filters at the transition of inlet to channel. Scale bar: 100 m. **(H)** Schematic showing the positioning of tissue sections compatible with xDbit. **(I)** Schematic cross section of a xDbit chip and clamp system.

**Figure S2.**
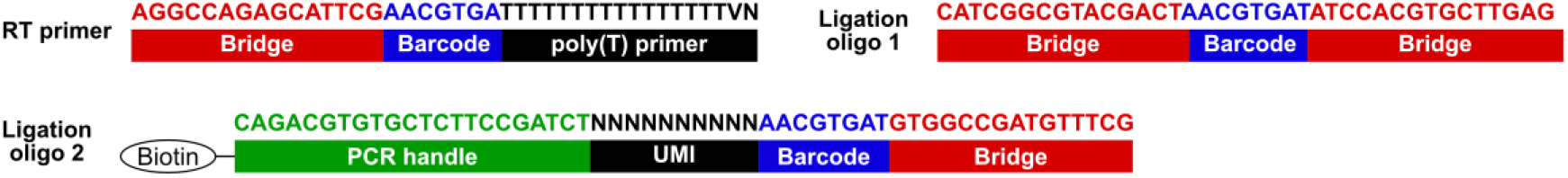
Sequences of barcoding oligonucleotides used during xDbit workflow. RevT: Reverse transcription; UMI: Unique molecular identifier; PCR: Polymerase chain reaction.

**Figure S3.**
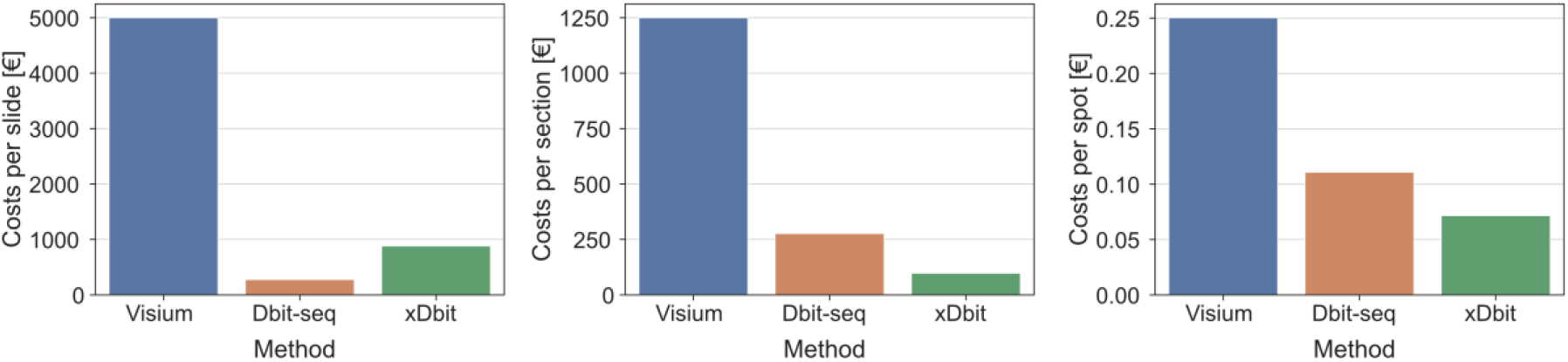
Comparison of xDbit costs with Visium Spatial (Salmén et al., 2018) and DBiT-seq method (Liu et al., 2020). The panels from left to right show the costs normalized per slide, section and spot, respectively. For DBiT-seq we calculated the costs from one experiment performed in our lab.

**Figure S4.**
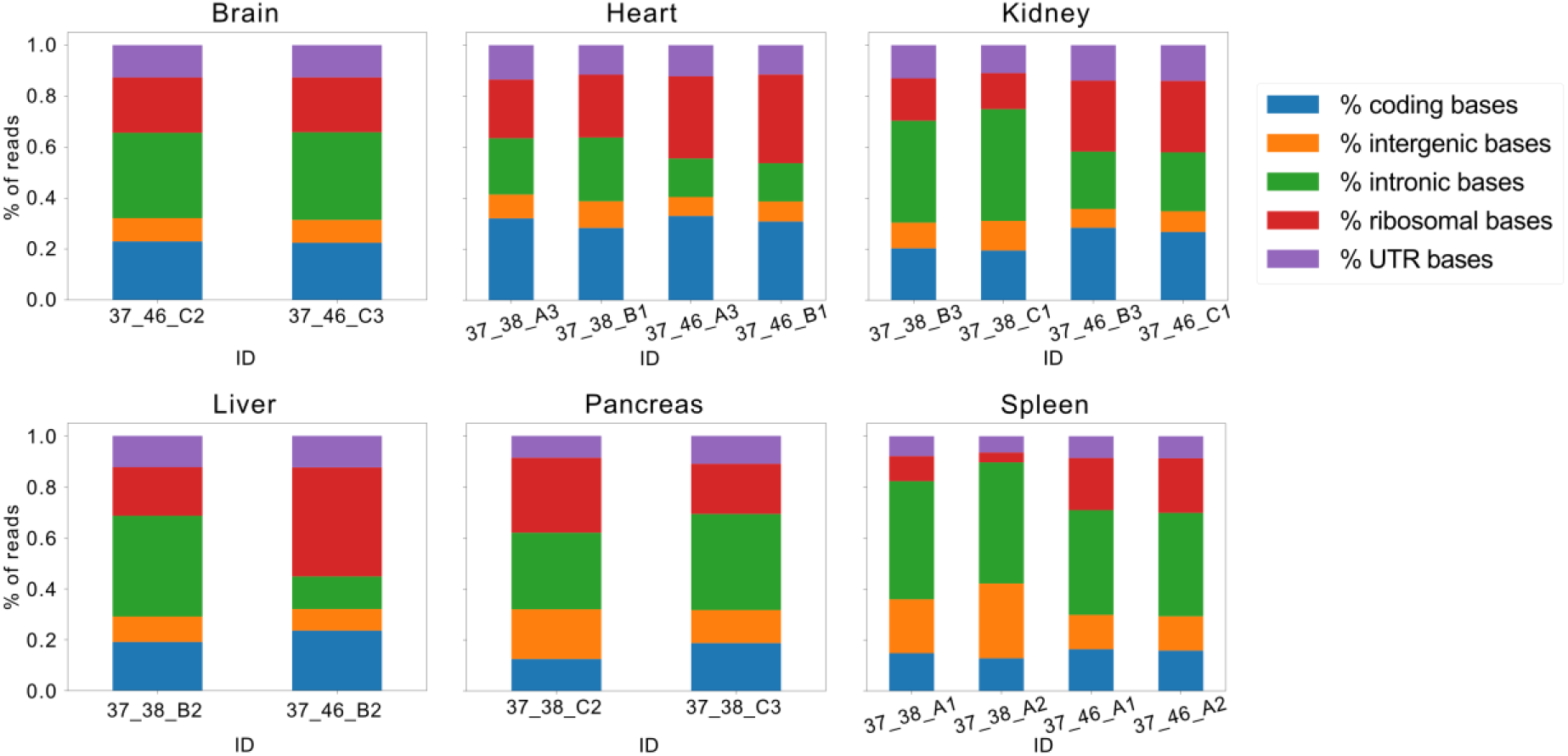
STAR alignment coverage metrics. Shown are the alignment coverage metrics for all experiments published in this study grouped by organ of origin.

**Figure S5.**
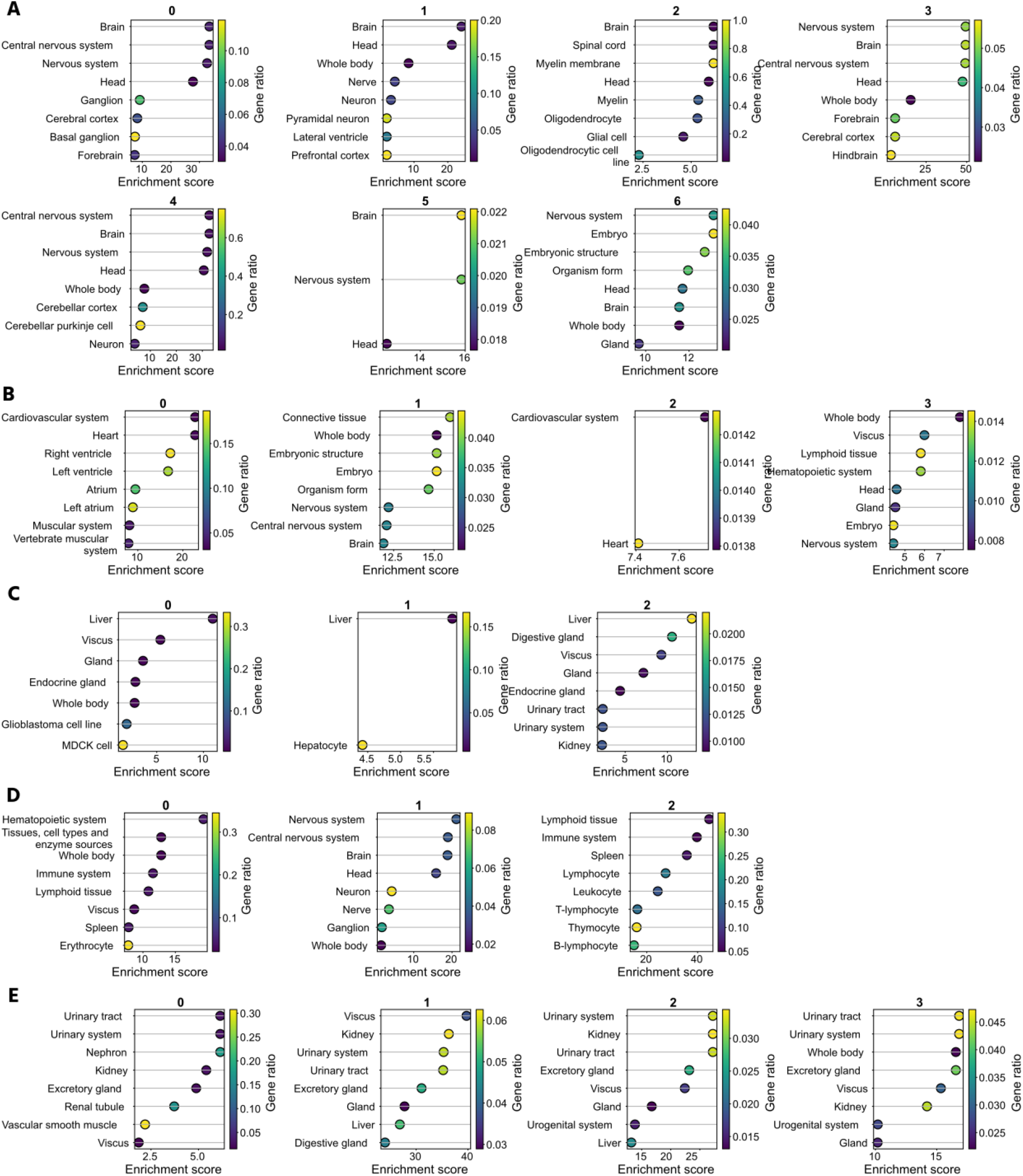
Gene ontology (GO) term enrichment analysis of *Leiden* clusters. GO term enrichment analysis was performed using the STRING algorithm (Szklarczyk et al., 2019) and the *Brenda Tissue Ontology* database (Gremse et al., 2011) for all Leiden clusters in cerebellum **(A)**, heart **(B)**, liver **(C)**, spleen **(D)**, and kidney **(E)**. The enrichment score was calculated as negative logarithmic p-value corrected for multiple testing using the Benjamini-Hochberg procedure. The gene ratio describes the size of the enrichment effect as logarithmic ratio of observed and expected number of genes.

**Figure S6.**
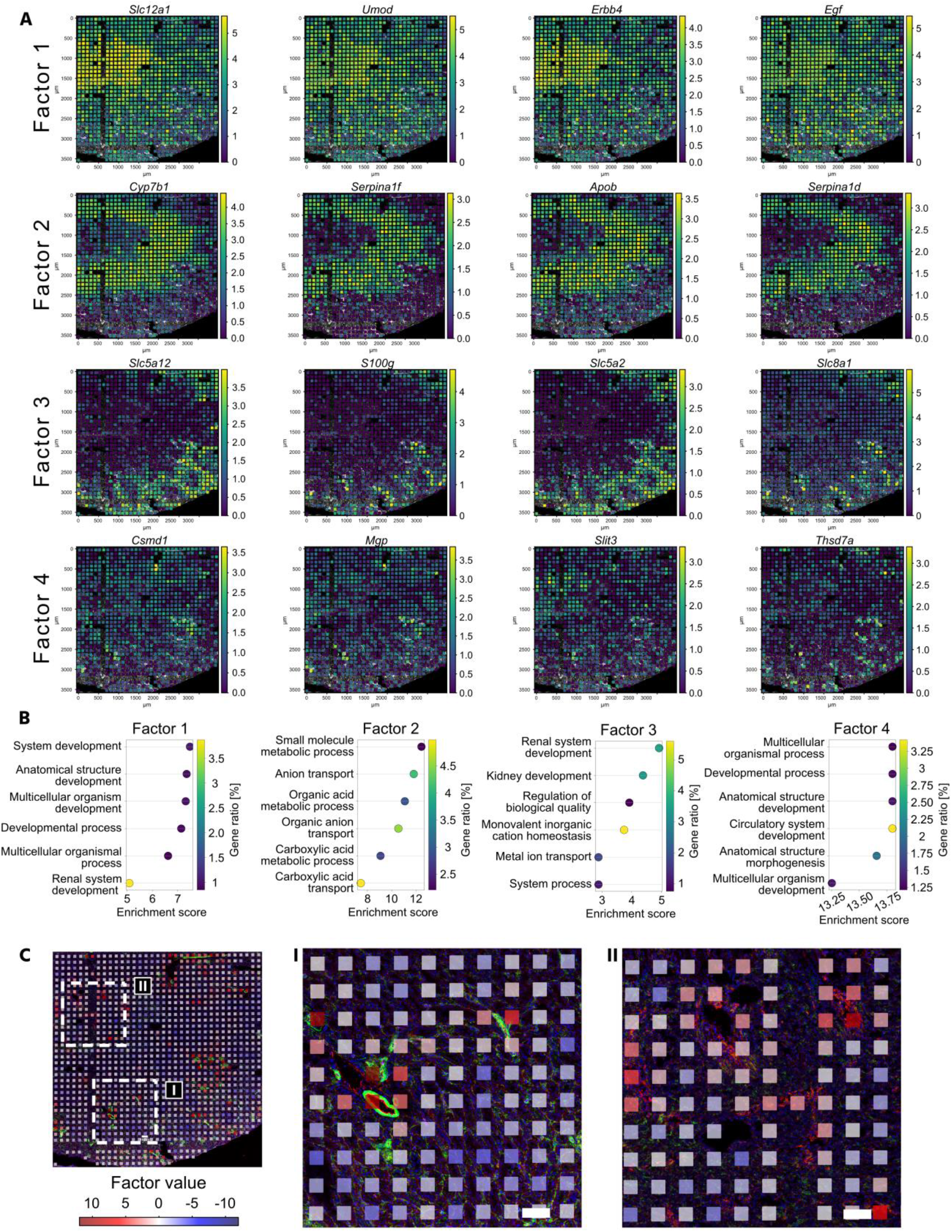
Extended factor analysis of one exemplary kidney section. Factor analysis was performed using MEFISTO (Velten et al., 2022). **(A)** Top positively weighted genes for the first 4 factors projected onto the respective phalloidin flourescence image. Colors denote the expression of the gene in the respective xDbit spot. **(B)** GO term enrichment analysis using the STRING algorithm (Szklarczyk et al., 2019) and the *Biological Processes* GO database (Ashburner et al., 2000; The Gene Ontology Consortium, 2021). As input we used the top positively weighted genes (> 95% CI) of the first four factors to reveal functional and structural areas of the murine kidney. CI: Confidence interval. **(C)** Details of the projection of MEFISTO factor 4 onto fluorescence images of the kidney section at two exemplary positions (I and II). Image colors denote CD31 (red), phalloidin (green) and DAPI (blue). The images show a correlation of the endothelial marker CD31 with factor 4. Scale bar: 100 μm.

**Figure S7.**
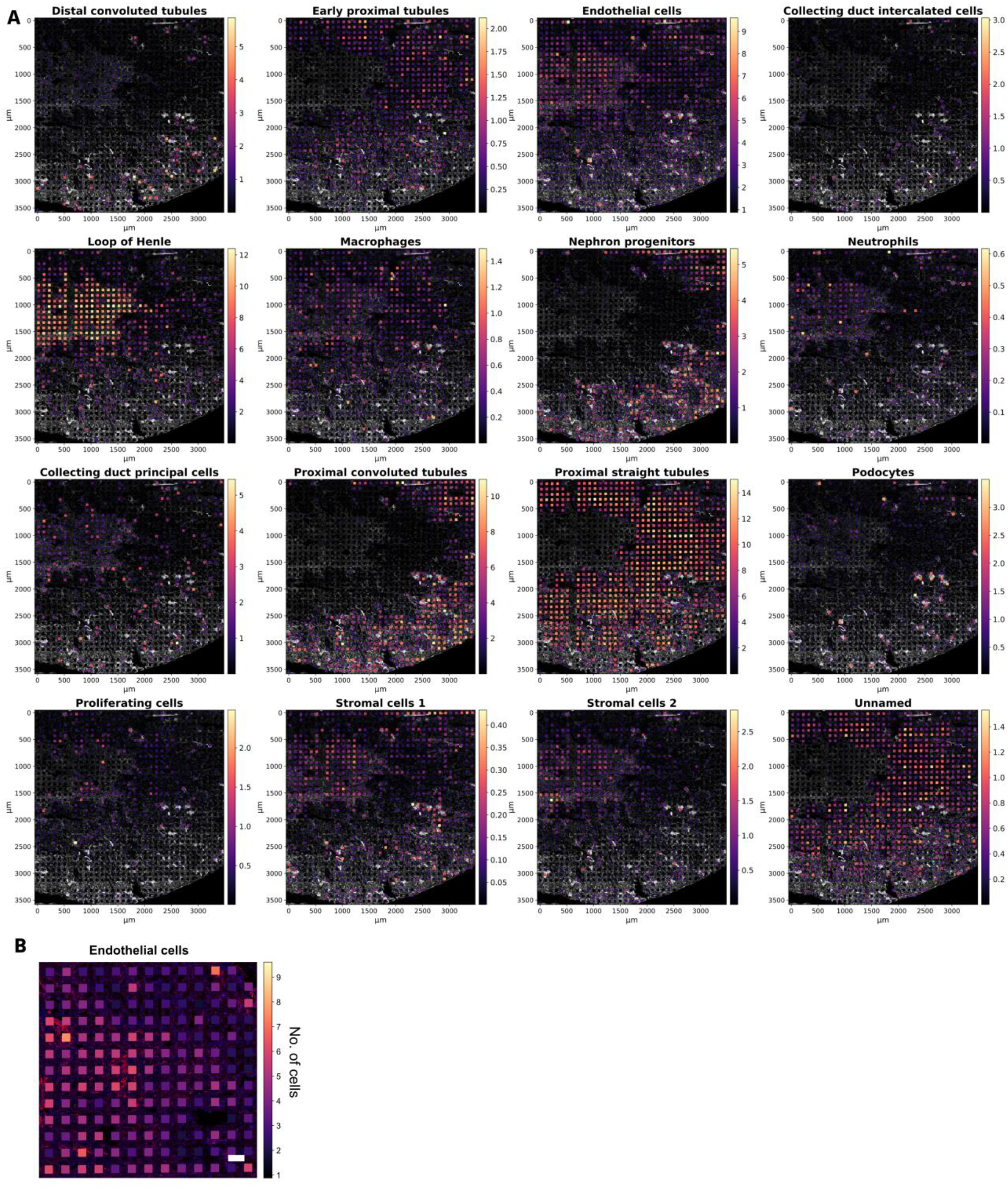
Extended cell type mapping results to the representative kidney section. Deconvolution of the xDbit ST results was performed using *cell2location* (Kleshchevnikov et al., 2022) and a published single-cell transcriptomics (scT) dataset of mural kidneys (Miao et al., 2021). **(A)** Spatially mapped proportions of all cell types, predicted by cell2location, onto the phalloidin fluorescence image of the kidney section. Spot colors denote the minimum number of cells predicted. **(B)** Detail of a representative region of the deconvolution results for endothelial cells. Spot colors denote the minimum number of predicted cells.

### Supplementary Tables

**Table S1.**
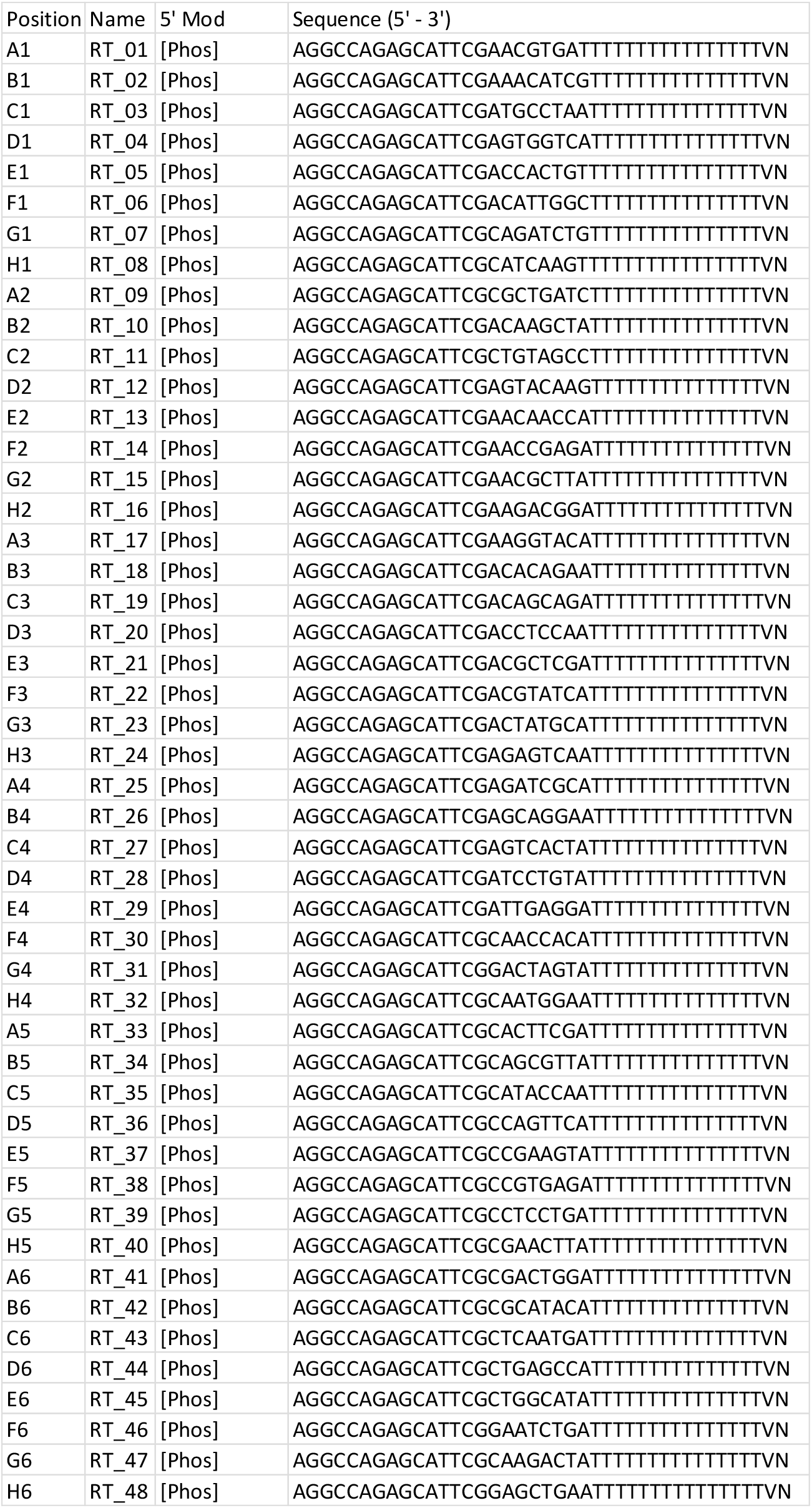
Sequences of primers for reverse transcription with respective 5’ modifications.

**Table S2.** Sets of RevT primer barcodes to encode the nine tissue sections. Optimal mixtures were determined using BARCOSEL (Somervuo et al., 2018).

|  |  |  |  |  |  |  |
| --- | --- | --- | --- | --- | --- | --- |
| Set #1 |  |  |  | Set #6 |  |  |
| Name | Sequence | Position |  | Name | Sequence | Position |
| >Round1_01 | AACGTGAT | A1 |  | >Round1_03 | ATGCCTAA | C1 |
| >Round1_27 | AGTCACTA | C4 |  | >Round1_06 | ACATTGGC | F1 |
| >Round1_44 | CTGAGCCA | D6 |  | >Round1_42 | CGCATACA | B6 |
| >Round1_46 | GAATCTGA | F6 |  | >Round1_47 | CAAGACTA | G6 |
| Set #2 |  |  |  | Set #7 |  |  |
| Name | Sequence | Position |  | Name | Sequence | Position |
| >Round1_05 | ACCACTGT | E1 |  | >Round1_29 | ATTGAGGA | E4 |
| >Round1_07 | CAGATCTG | G1 |  | >Round1_33 | CACTTCGA | A5 |
| >Round1_11 | CTGTAGCC | C2 |  | >Round1_38 | CCGTGAGA | F5 |
| >Round1_24 | AGAGTCAA | H3 |  | >Round1_40 | CGAACTTA | H5 |
| Set #3 |  |  |  | Set #8 |  |  |
| Name | Sequence | Position |  | Name | Sequence | Position |
| >Round1_02 | AAACATCG | B1 |  | >Round1_17 | AAGGTACA | A3 |
| >Round1_09 | CGCTGATC | A2 |  | >Round1_25 | AGATCGCA | A4 |
| >Round1_45 | CTGGCATA | E6 |  | >Round1_34 | CAGCGTTA | B5 |
| >Round1_48 | GAGCTGAA | H6 |  | >Round1_43 | CTCAATGA | C6 |
| Set #4 |  |  |  | Set #9 |  |  |
| Name | Sequence | Position |  | Name | Sequence | Position |
| >Round1_12 | AGTACAAG | D2 |  | >Round1_15 | AACGCTTA | G2 |
| >Round1_21 | ACGCTCGA | E3 |  | >Round1_22 | ACGTATCA | F3 |
| >Round1_31 | GACTAGTA | G4 |  | >Round1_26 | AGCAGGAA | B4 |
| >Round1_36 | CCAGTTCA | D5 |  | >Round1_41 | CGACTGGA | A6 |
| Set #5 |  |  |  |  |  |  |
| Name | Sequence | Position |  |  |  |  |
| >Round1_04 | AGTGGTCA | D1 |  |  |  |  |
| >Round1_08 | CATCAAGT | H1 |  |  |  |  |
| >Round1_20 | ACCTCCAA | D3 |  |  |  |  |
| >Round1_28 | ATCCTGTA | D4 |  |  |  |  |

**Table S3.**
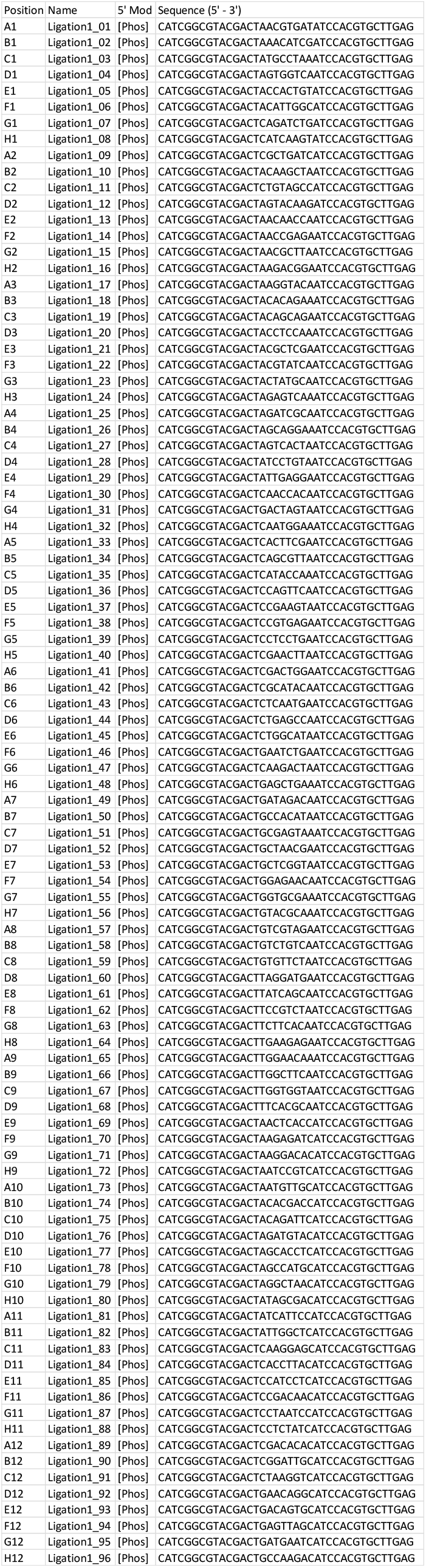
Sequences for ligation round 1 oligonucleotides and respective 5’ modifications.

**Table S4.**
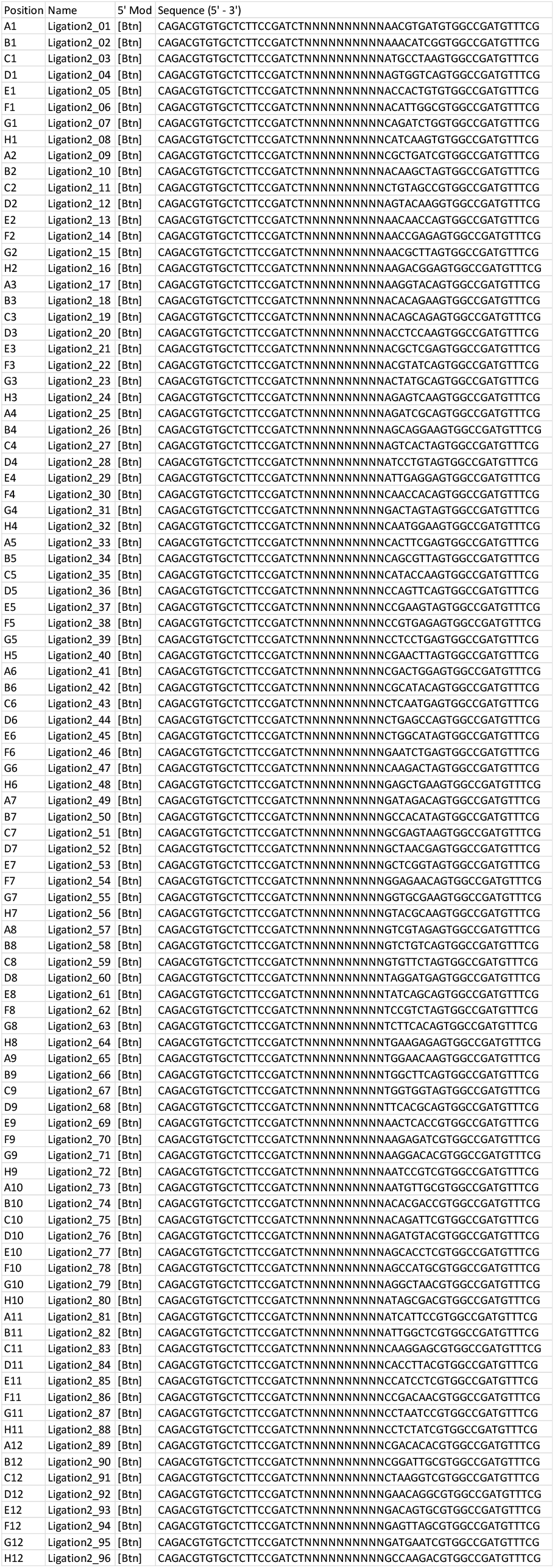
Sequences for ligation round 2 oligonucleotides and respective 5’ modifications.

**Table S5.**
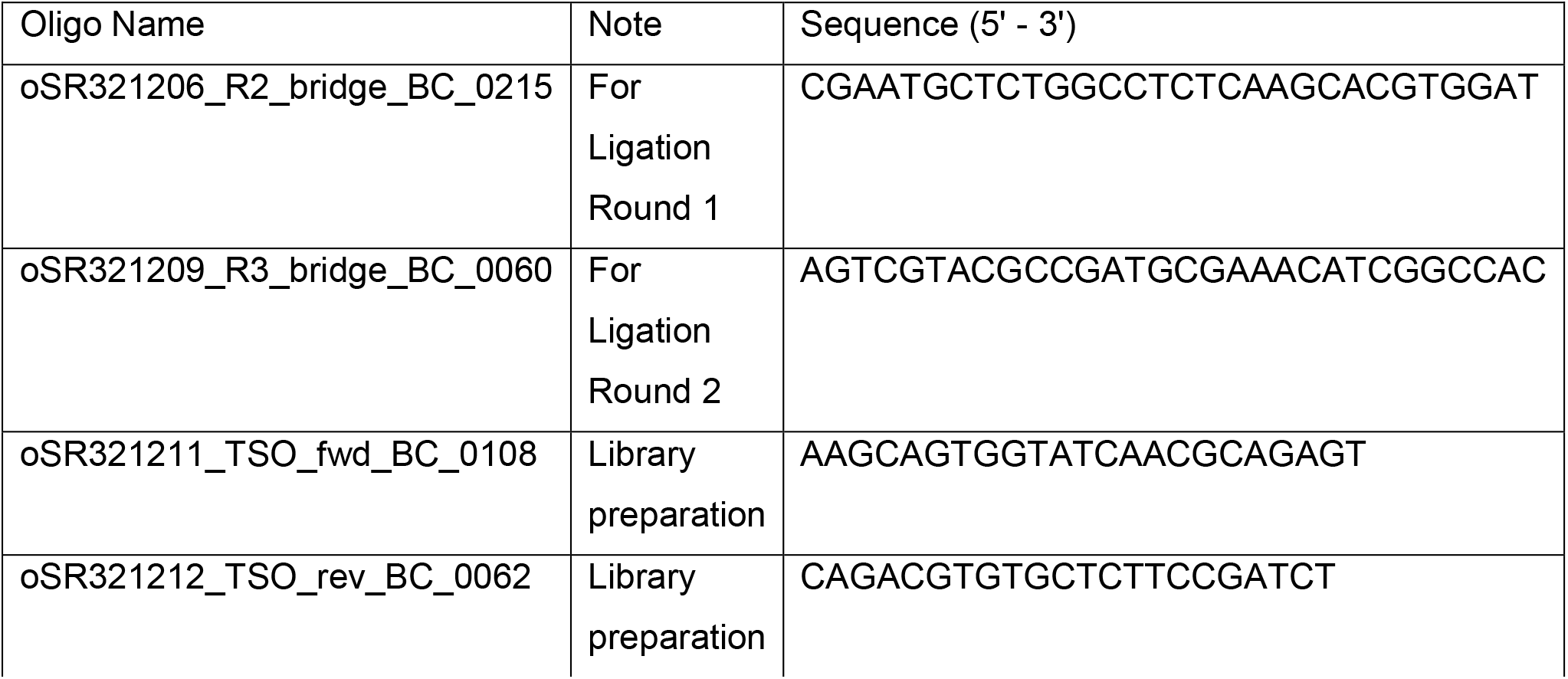
Sequences of the short bridge oligonucleotides and primers used during xDbit library preparation.

**Table S6.** Filling scheme of inlets for barcoding rounds with the xDbit microfluidic chip. Inlets filled with alignment markers and ligation barcodes are colored red and green, respectively.

|  | 1 | 2 | 3 | 4 | 5 |  |  |
| --- | --- | --- | --- | --- | --- | --- | --- |
| A |  | 9 | 17 | 25 |  |  | Alignment marker |
| B | 2 | 10 | 18 | 26 | 34 |  | Barcode # |
| C | 3 | 11 | 19 | 27 | 35 |  |  |
| D | 4 | 12 | 20 | 28 | 36 |  |  |
| E | 5 | 13 | 21 | 29 | 37 |  |  |
| F | 6 | 14 | 22 | 30 | 38 |  |  |
| G | 7 | 15 | 23 | 31 | 39 |  |  |
| H |  | 16 | 24 | 32 |  |  |  |

**Table S7.** Cell type percentages obtained from the cell2location analysis and subsequential comparsion to published percentages from Clark et al., 2019. Medullary thick ascending loop (mTAL), principal cells (PCs), Cortical collecting duct (CCD), Outer medullary collecting duct (OMCD), Intercalated cells type A and B (A-ICs + B-ICs).

| Cell type | Cell2location [%] | Clark et al., 2019 [%] |
| --- | --- | --- |
| Distal convoluted tubules | 1.91 | 7.45 |
| Early proximal tubules | 2.29 |  |
| Endothelial cells | 17.27 |  |
| Collecting duct intercalated cells | 0.68 | 6.96 (All A-ICs + B-ICs) |
| Loop of Henle | 15.70 | 14.87 (mTAL) |
| Macrophages | 1.54 |  |
| Nephron progenitors | 4.40 |  |
| Neutrophils | 0.38 |  |
| Collecting duct principal cells | 2.87 | 4.96 (CCD PC + OMCD PC) |
| Proximal convoluted tubules | 10.77 | 44 |
| Proximal straight tubules | 34.55 |  |
| Podocytes | 1.67 | 3 |
| Proliferating cells | 1.12 |  |
| Stromal cells 1 | 0.47 |  |
| Stromal cells 2 | 2.01 |  |
| Unnamed | 2.37 |  |

## Notes

https://github.com/jwrth/xDbit_toolbox

https://www.ncbi.nlm.nih.gov/geo/query/acc.cgi?acc=GSE207843

### Competing Interest Statement

The authors have declared no competing interest.

